# Prefrontal gamma oscillations engage dynamic cell type-specific configurations to support flexible behavior

**DOI:** 10.1101/2024.03.08.584173

**Authors:** Aarron J. Phensy, Lara L. Hagopian, Caitriona M. Costello, Simon Haziza, Omkar Ghenand, Yanping Zhang, Mark J. Schnitzer, Vikaas S. Sohal

**Affiliations:** Department of Psychiatry and Behavioral Sciences, San Francisco, CA 94143-0444; Weill Institute for Neurosciences, Center for Integrative Neuroscience, and Kavli Institute for Fundamental Neuroscience, University of California, San Francisco, San Francisco, CA 94143-0444; James H. Clark Center, Stanford University, Stanford, CA 94305, USA; CNC Program, Stanford University, Stanford, CA 94305, USA; Department of Biology, Stanford University, Stanford CA 94305, USA; Howard Hughes Medical Institute, Stanford University, Stanford, CA 94305, USA; Department of Applied Physics, Stanford University, Stanford CA 94305, USA; Department of Neurosurgery, Stanford University, Stanford CA 94305, USA

## Abstract

Cognitive dysfunction in conditions such as schizophrenia involves disrupted communication between the prefrontal cortex (PFC) and mediodorsal thalamus (MD). Parvalbumin interneurons (PVI) are known to regulate PFC microcircuits and generate gamma-frequency (∼40Hz) oscillations – fast, synchronized neural rhythms that are recruited during many executive functions, necessary for cognitive flexibility, and deficient in schizophrenia. While targeting PVI-mediated gamma oscillations holds great therapeutic promise, their nature and specific functions, e.g., for regulating PFC→MD communication, remain elusive. Using dual-color voltage indicators and optogenetics, we reveal that PVIs dynamically entrain MD-projecting PFC neurons both locally and contralaterally, giving rise to multiple distinct circuit-specific patterns of distributed synchronization that are recruited in a behaviorally-specific manner to support particular aspects of flexible behavior. Thus, gamma oscillations are not unitary phenomena characterized by one microcircuit-wide pattern of entrainment. Rather, they comprise diverse motifs, defined by specific cell types and phase relationships, that are dynamically recruited for specific functions.

## INTRODUCTION

Cognitive deficits are a core symptom of schizophrenia that profoundly impact quality of life for affected individuals^1,2^. Many of these deficits are attributed to dysfunction involving neural circuits within the prefrontal cortex^3^ as well as its interactions with other regions such as the mediodorsal thalamus^4,5^ and dorsal striatum^6,7^. Of note, deficits in neural oscillations in the gamma frequency band (∼30-100 Hz) are heavily documented in schizophrenia, including a reduction in evoked gamma oscillatory power during tasks which involve cognitive control and flexible behavior such as the Wisconsin Card Sorting Task (WCST)^3,8^. This has led to substantial interest in targeting prefrontal gamma oscillations and their underlying mechanisms in order to enhance cognitive function in schizophrenia as well as other neuropsychiatric conditions^9–11^. However, this requires a deeper understanding of the precise nature, underlying mechanisms, and relevant functions of gamma oscillations ^12–14^.

Parvalbumin-expressing GABAergic interneurons (PVI) comprise a substantial fraction of all cortical interneurons and are believed to be a critical mediator of gamma oscillations in the brain^15^. Importantly, reductions in markers of PVI function are one of the core pathologies associated with schizophrenia^16^ and likely a major contributor to the altered gamma activity observed in individuals with schizophrenia. In animals, loss of PVI function in the medial prefrontal cortex (mPFC) has extensively been studied^17,18^ and results in aberrant gamma oscillatory activity^18,19^ and impaired cognitive flexibility^20,21^. Recent work from our lab has identified that prefrontal PVIs not only generate local gamma oscillations, but also synchronize gamma activity across the left and right prefrontal cortices as mice perform a task involving cognitive flexibility^22^. Importantly, by comparing the effects of 40Hz optogenetic stimulation delivered either in-phase or out-of-phase across the hemispheres, these studies confirmed that this cross-hemispheric gamma synchrony between prefrontal PVIs (‘PVI-xH-γ-sync’) is *necessary* for mice to appropriately perform the task.

Furthermore, optogenetically restoring PVI-xH-γ-sync can rescue task performance in mutant mice with impaired PVI function^22^. Importantly, PVIs are known to entrain deep layer projection neurons through a combination of feedforward and feedback inhibition^15,23^. More recently, we have found that parvalbumin-expressing inhibitory neurons also target deep layer projection neurons (particularly those projecting to the mediodorsal thalamus) in the contralateral mPFC, and that these callosal PV+ projections play essential roles in gamma synchronization and cognitive flexibility^24^.

Thus cross-hemispheric PVI gamma synchrony plays a central role in cognitive flexibility, and PVIs are well-positioned to propagate their synchrony to deep layer projection neurons, including those targeting mediodorsal thalamus. This raises two crucial questions. First, do PVIs actually propagate synchrony to other cell types? Second, can we identify the specific aspects of cognitive flexibility to which this synchrony contributes (e.g., learning from trial outcomes, decision making, cognitive control, etc.)? Answering these questions is critical for understanding the function of gamma oscillations in normal cognition and identifying specific features of gamma oscillations that mediate their therapeutic effects. Here, using genetically encoded voltage indicators and optogenetics, we confirm that indeed, PVIs selectively entrain deep layer neurons which project to the MD thalamus. However, surprisingly, this transmission of gamma synchrony only occurs at specific behavioral timepoints and does not globally entrain other neuronal populations, as we did not observe similar patterns of synchronization for projection neurons targeting the dorsal striatum. Further, we find that PVIs form unique phase configurations with both local and contralateral MD-projection neurons that evolve across learning in a context-specific manner. Thus, gamma oscillations are not a singular entity, but actually comprise multiple distinct synchrony configurations, each of which may enable unique cortical computations that can enable specific behavioral outputs.

## RESULTS

### Mice transition between stable strategies when adapting to changes in task structure

In order to examine the neural mechanisms underlying cognitive flexibility, we utilized a variant of the bowl-digging rule-shift task which has been used extensively in our lab^18^. In this task (Fig. 1A-B), on each trial mice are presented with two bowls and must correctly choose one to obtain a food reward. Bowls contain compound cues, i.e., each bowl contains 1 of 4 possible odor + texture pairings, either garlic + sand, coriander + litter, garlic + litter, or coriander + sand. Mice have to first learn to dig in the bowl identified by a single cue from one modality. For example, during the ‘Initial Association’ (IA) phase, the mice might learn to dig in the *sand*-containing bowl. This would be an example of a texture rule for which mice can simply ignore odor cues. Then, during the ‘Rule Shift’ (RS) phase, they must learn to associate reward with a different cue belonging to the previously ignored modality, e.g., by digging in the *garlic*-containing bowl. This would be an example of an odor rule for which mice can simply ignore texture cues. This extradimensional shift across modalities requires a functioning PFC^25^, is dependent on PVI-activity^18^, and takes significantly more trials to learn than the initial association (t_(56)_ = 6.13, p < 0.001; Fig. 1C). Importantly, to enhance our ability to time-lock neural recordings to discrete behavioral moments we implemented a change from our previously-used behavioral protocol: here, instead of pre-baiting each correct bowl with a hidden food reward, when a mouse dug in their choice bowl, the experimenter either provided a reward with forceps (correct choice) or removed the correct bowl (error choice), thus providing precisely-timed and cued outcomes (Fig. 1A). Interestingly, this extra step slightly increased trials to criterion compared to our prior studies, which allowed us to examine learning across a larger spread of trials.

**Figure 1.**
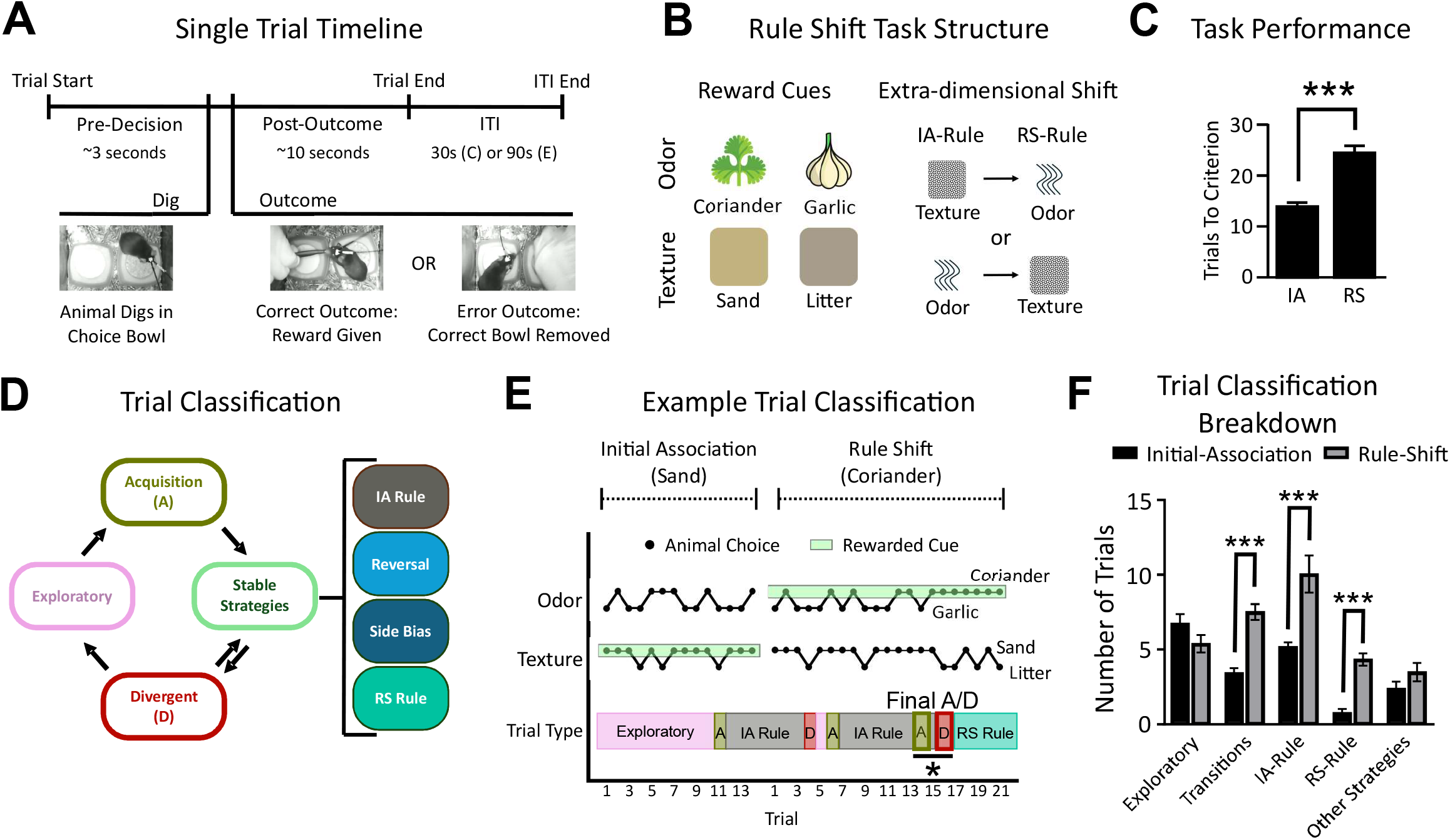
Behavioral framework reveals dynamic strategy transitions during rule shift learning. (A) Schematic of single trial timeline in the extra-dimensional rule shift task. Mice initiate a trial and choose between two bowls containing compound odor-texture cues. Once mice dig in a bowl, the experimenter either provides a food reward (correct trial) or removes the correct bowl (error trial), providing a time-locked outcome signal. (B) Examples of compound cues and rule shift design. Mice first learn an Initial Association (IA) rule (e.g., dig in bowls containing a specific texture), then perform a Rule Shift (RS) requiring them to shift to a different rewarded modality (e.g., odor). (C) Mice require significantly more trials to reach criterion during RS compared to IA (***p < 0.001). (D) Schematic of behavioral strategy classification using a machine learning algorithm trained on past and future trial predictability. The behavioral strategy on each trial is categorized as Stable, Exploratory, Divergent (D), Acquisition (A), or bias-driven (e.g., side bias). (E) Example of dynamic trial classification during learning, showing transitions between strategies as the mouse progresses through learning of the IA and RS. (F) Quantification of trial types shows that during RS, mice exhibit more transitions between stable strategies and more Acquisition/Divergent trials than during IA (***p < 0.001).

We observed that mice did not consistently follow a single, predictable learning trajectory. Rather, they appeared to utilize a combination of perseverative / regressive strategies, random choices, and other strategies such as side biases or reversals. To tease apart these different strategies, we designed a machine learning algorithm that classified trials based on the predictability of the mouse’s choice given past and future trials (Fig. 1D-E). This allowed us to categorize behavior into blocks in which the algorithm could (stable strategies) or could not (exploratory behavior) successfully predict the animal’s choice. Divergent trials mark departures from a stable choice policy, while acquisition trials correspond to the onset of a new stable policy. Given the learning criterion (8 correct out of 10 consecutive trials) in both IA and RS phases, behavior is structured into stable blocks, contributing to the robustness of classification metrics across window sizes (Supp. Fig. 1F–I). Interestingly, when we compared the rule-shift portion of the task to the initial-association portion, we found a significant increase in the number of transition and stable policy trials – revealing that during rule shifts, mice spend more time transitioning between stable strategies as opposed to simply following exploratory or unpredictable choice selection strategies (Two-way ANOVA: Task Phase: F_(1,560)_ = 41.93, p < 0.001; Trial Class: F_(4,560)_ = 25.65, p < 0.001; Interaction: F_(4,560)_ = 9.06, p < 0.001; Fig. 1F).

### PFC PVIs increase γ-synchrony immediately after trial outcomes when mice change behavioral strategies

Our lab previously found that PVIs synchronize at gamma frequencies (30-50 Hz) across the left and right prefrontal cortices during rule-shifts – specifically following error trials^22^. However, this analysis measured synchrony over timescales of minutes, making it unclear whether increases in PVI-xH-γ-sync begin immediately (within seconds) following errors or later (e.g., during the subsequent inter-trial interval). To better understand the temporal dynamics of PVI-xH-γ-sync, we expressed the genetically encoded voltage indicator (GEVI) Ace2N-4AA-mNeon (Ace-mNeon) in prefrontal PVIs (Fig. 2A-B) and performed TEMPO (trans-membrane electrical measurements performed optically)^22,26^, which allows for cell type-specific measurements of voltage activity with high temporal resolution sufficient to track gamma-frequency oscillations (∼30–50 Hz). We then developed a novel analysis pipeline that enabled the precise measurement of interhemispheric synchrony during epochs as brief as 1s (Fig. 2C, Methods). To confirm that our synchrony measurements reflect true phase relationships rather than shared power or artifacts, we validated the pipeline three ways: by 1) analyzing simulated data with known synchronization, 2) using phase-scrambling to disrupt synchronization in real data, and 3) calculating synchrony between voltage signals and the reference channel. These controls validated that our method only detected genuine synchrony driven by voltage signals (Supp. Fig. 2). We used this pipeline to examine synchrony between left and right prefrontal PVIs during the pre-decision and post-outcome periods across four trial types defined within the **Rule Shift (RS) phase** (Fig. 1), categorized as either **Stable Strategies** (*IA-Strategy, RS-Strategy*) or **Behavioral Transitions** (*IA-Divergent, RS-Acquisition*). *IA-Strategy* trials reflect continued use of the outdated Initial Association rule, while *RS-Strategy* trials reflect stable application of the newly learned RS rule. *IA-Divergent* and *RS-Acquisition* trials mark the final deviation from IA-based behavior and the final error before strategy stabilization, respectively— timepoints which represent key learning events. These trial types were selected to focus analyses on behaviorally meaningful moments of strategy execution and updating. Interestingly, we did not observe any changes in PVI γ-synchrony during the pre-decision period for any of the four trial types (F_(3,45)_ = 1.02, p = 0.39; Fig. 2D). However, PVI-xH-γ-sync did increase during the post-outcome period following both IA-Divergent and RS-Acquisition trials (F_(3,45)_ = 6.15, p = 0.001; Fig. 2E). Importantly, these two trial types represent periods of transition during which the mouse is in the process of updating their choice behavior, suggesting that prefrontal PVI synchrony is associated with updating behavioral strategies.

**Figure 2.**
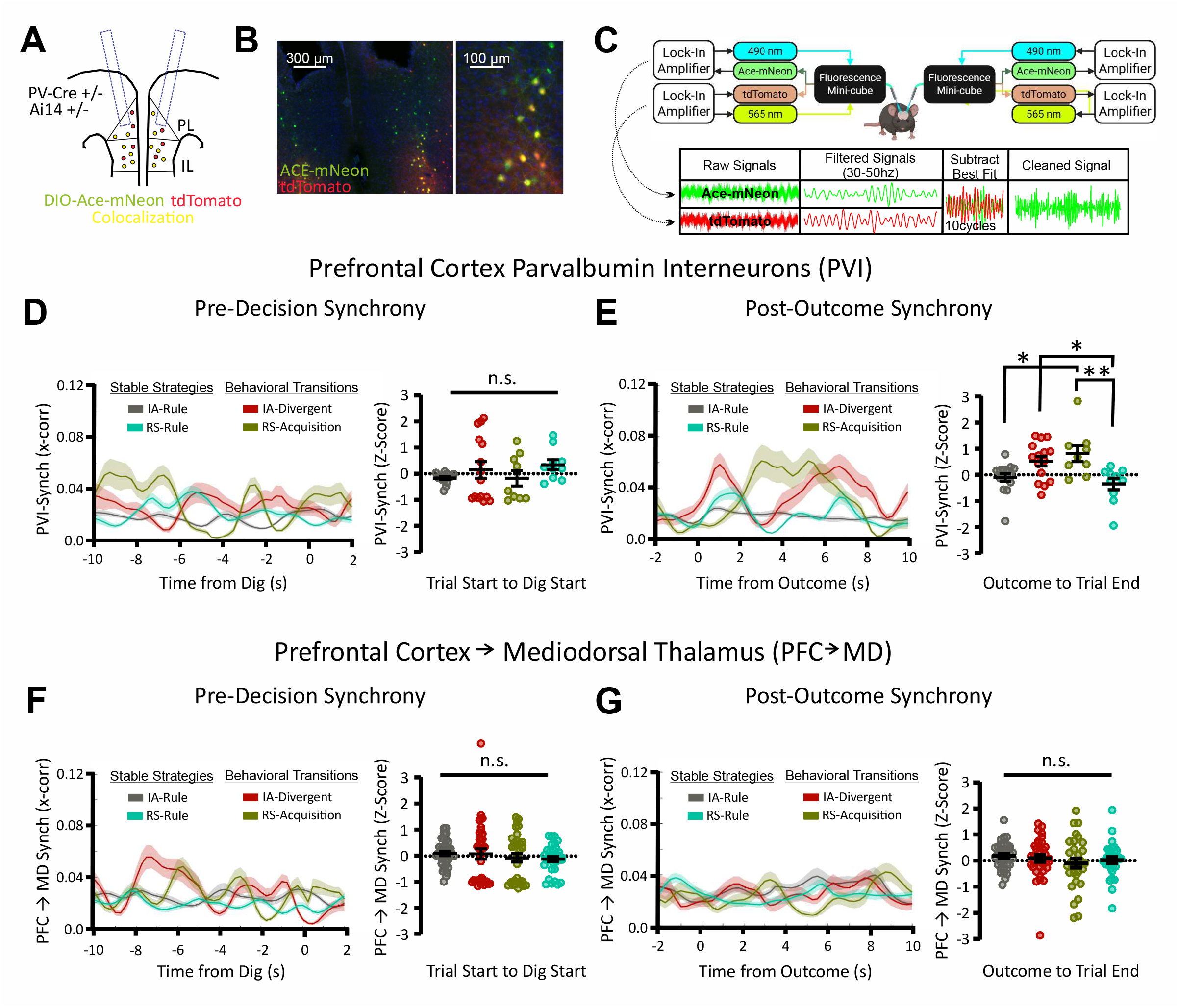
Interhemispheric gamma synchrony between prefrontal PV interneurons increases specifically after behavioral transitions. (A) Schematic showing bilateral injections of AAV-DIO-Ace2N-4AA-mNeon (Ace-mNeon) into the medial prefrontal cortex (mPFC) of PV-Cre+/−; Ai14+/− mice to label PV interneurons (PVIs). PL: prelimbic cortex; IL: infralimbic cortex. (B) Histological validation showing Ace-mNeon expression in PV interneurons (green) and Ai14-driven tdTomato labeling (red). (C) Schematic of the hardware and analysis workflow. Dual-site fiber photometry utilizing lock-in amplifier demodulation to separate Ace-mNeon and tdTomato fluorescence signals. Fluorescence signals subsequently undergo regression (to remove artifacts) and filtering (to isolate gamma-band activity). Resulting voltage signals are used for cross-correlation analysis. (D) Left: Example time course of gamma synchrony (cross-correlation of gamma-band filtered signals from left and right hemispheres), aligned to dig onset, to illustrate signal dynamics across trial types. Right: Quantification of gamma synchrony during the pre-decision window, normalized to a pre-task baseline, showed no significant modulation across trial types. (E) After trial outcomes, gamma synchrony increased specifically on trials corresponding to behavioral transitions (the IA-Divergent and RS-Acquisition trials) compared to trials associated with stable behavioral strategies (*p<0.05, **p<0.01), indicating that post-outcome gamma synchrony is recruited when mice update their strategy. (F) Similar to D, but for interhemispheric gamma synchrony between mPFC neurons projecting to the mediodorsal thalamus (PFC→MD). PFC→MD neurons were labeled using retrograde CAV2-Cre injections into MD. Pre-decision gamma synchrony in PFC→MD neurons showed no differences across trial types. (G) Similar to E, but for interhemispheric gamma synchrony between PFC→MD neurons. In contrast to PVIs, no significant changes in post-outcome gamma synchrony were observed for PFC→MD neurons during the post-outcome periods of trials corresponding to behavioral transitions.

Gamma oscillations are believed to reflect rhythmic fluctuations in perisomatic inhibition, originating mainly from PVIs, onto pyramidal neurons^27,28^. In this way, rhythmic activity in PVIs is believed to entrain excitatory neurons, suggesting that increased gamma synchrony between PVIs in the left and right mPFC may also be transmitted to projection neurons. To examine this, we expressed Ace-mNeon and tdTomato in two populations of prefrontal projection neurons believed to play important roles in cognitive flexibility: deep layer projection neurons targeting either the mediodorsal thalamus (PFC→MD) or the dorsal striatum (PFC→DS). To our surprise, neither PFC→MD neurons (Pre-Decision: F_(3,132)_ = 0.54, p = 0.65; Fig. 2F. Post-Outcome: F_(3,132)_ = 0.81, p = 0.49; Fig. 2G) nor PFC→DS neurons (Pre-Decision: F_(3,32)_ = 0.24, p = 0.87; Post-Outcome: F_(3,32)_ = 0.6, p = 0.62; data not shown) exhibited an increase in interhemispheric gamma synchrony during the post-outcome period on IA-Divergent or RS-Acquisition trials. Thus, the increase in PVI interhemispheric gamma synchrony observed as mice shift behavioral strategies is not automatically transmitted across local microcircuits into the neurons which are major recipients of PVI-mediated perisomatic inhibition.

### Changes in PFC→MD neuron γ-synchrony track deviation from the IA strategy

The preceding observation made us wonder whether there are any times when we could detect increased cross-hemispheric gamma synchrony for specific populations of projection neurons, or whether such epochs of elevated synchrony might never occur (due to the nature of these cells and/or their connectivity), or might not be able to be resolved (due to limitations of TEMPO in these neurons). Importantly, it has been shown that PFC→MD projections influence choice selection^29^, therefore we were particularly interested in whether changes in gamma synchrony involving PFC→MD projection neurons might occur during the pre-decision period, when mice choose which bowl to dig in. Notably, rule-shift period can be divided into two types based on the difficulty of making this decision. In half of trials (“non-conflict” trials), the cue that was previously rewarded during the Initial Association and the cue that becomes rewarded during the Rule Shift cue are both located in the same bowl. Thus, mice would make a correct choice by following either rule. Conversely, in the other half of trials (“conflict” trials), the Initial Association and Rule Shift cues are located in different bowls, such that selecting the bowl containing the cue that was rewarded during the Initial Association would result in a perseverative error (Fig. 3A-B). Indeed, mice perform significantly better on non-conflict than conflict trials, reflecting their perseverative behavior (Two-way ANOVA: non-conflict vs conflict: F_(1,224)_ = 0.004, p = 0.95; correct vs error: F_(1,224)_ = 17.97, p < 0.001; Interaction: F_(1,224)_ = 52.53, p < 0.001; Fig. 3C).

**Figure 3.**
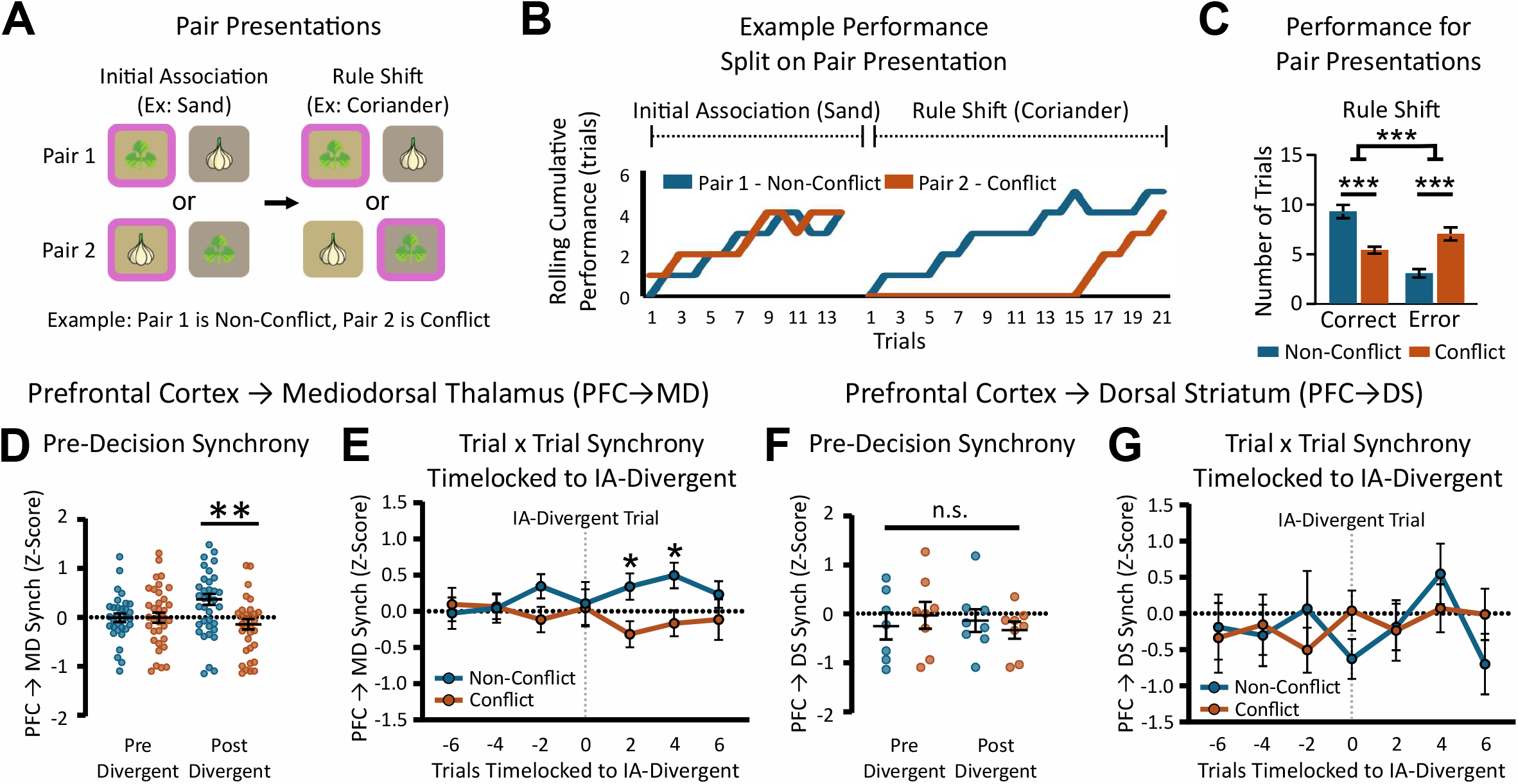
Conflict-dependent modulation of gamma synchrony in PFC→MD neurons during decision-making. (A) Schematic illustrating how rule-shift trials are categorized as *conflict* or *non-conflict* depending on whether the Initial Association (IA) and Rule Shift (RS) cues are present in the same or different bowls, respectively. On non-conflict trials, following either the old (IA) or new (RS) rule results in a correct choice, whereas conflict trials require ignoring the outdated IA rule to choose correctly. (B) Performance of an example mouse across conflict and non-conflict trials, showing that mice perform worse on conflict trials, reflecting perseveration. (C) Quantification of correct and error choices shows significantly lower accuracy on conflict trials compared to non-conflict trials (***p<0.001). Notably, higher correct choices on non-conflict trials and higher errors on conflict trials both reflect continued use of the outdated IA strategy. (D) After the IA-Divergent trial, interhemispheric gamma synchrony between PFC→MD neurons during the pre-decision period differs depending on trial type: it increases on non-conflict trials but decreases slightly on conflict trials. (E) Trial-by-trial analysis of pre-decision interhemispheric gamma synchrony between PFC→MD neurons shows that this difference (higher synchrony on non-conflict vs. conflict trials) emerges 2-4 trials after the IA-Divergent trial. (F) Pre-decision interhemispheric gamma synchrony between PFC→dorsal striaum (DS) projection neurons does not differ on conflict vs. non-conflict trials before or after the IA-Divergent trial. (G) Similar to E but for pre-decision interhemispheric gamma synchrony between PFC→DS neurons. Trial-by-trial analysis of PFC→DS neuron synchrony showed no modulation related to conflict or behavioral transitions.

We examined interhemispheric gamma synchrony for either PFC→MD or PFC→DS neurons during the pre-decision period as a function of trial type (non-conflict vs. conflict) as mice progressed through the rule-shift task. Crucially, the non-conflict vs. conflict distinction is behaviorally irrelevant during the initial association. Thus, differences in neural activity during the pre-decision period between conflict and non-conflict trials should only appear when the nature of the rule transition becomes apparent to mice. Indeed, interhemispheric gamma synchrony between PFC→MD neurons in the left and right mPFC did not differ between the pre-decision periods of conflict and non-conflict trials prior to the IA-Divergent trial – the trial on which mice finally stop following the IA rule. Following the IA-Divergent trial, PFC→MD neurons began to exhibit significantly higher interhemispheric gamma synchrony during pre-decision periods on non-conflict trials than conflict trials (Mixed Effects Model: Trial Type: F_(1,34)_ = 3.73, p = 0.06; Divergent Split: F_(1,34)_ = 0.05, p = 0.83; Interaction: F_(1,28)_ = 8.12, p = 0.008; Fig. 3D). Examining synchrony on a trial-by-trial basis revealed that this effect was primarily driven by the first 4 trials following the IA-Divergent trial (Mixed Effects Model: Trial : F_(4.8,217.9)_ = 0.19, p = 0.96; Trial Type: F_(1,68)_ = 4.354, p = 0.04; Fig. 3E). Interestingly, when we examined PFC→DS neurons we did not observe any differences in their interhemispheric gamma synchrony based on trial type (Two-way ANOVA: Trial Type: F_(1,27)_ = 0.002, p = 0.96; Divergent Split: F_(1,27)_ = 0.16, p = 0.69; Interaction: F_(1,27)_ = 0.75, p = 0.51; Fig. 3F) nor using a trial-by-trial analysis (Mixed Effects Model: Trial #: F_(3.5,28.3)_ = 0.73, p = 0.57; Trial Type: F_(1,14)_ = 0.04, p = 0.85; Fig. 3G). Thus, changes in PFC→MD neuron gamma synchrony encode task features that only become relevant after the rule transition and align with changes in behavior, specifically in the strategies that mice use to make choices.

### PVIs are necessary for changes in PFC→MD neuron γ-synchrony

We did not observe an increase in PVI-xH-γ-sync during the pre-decision period. This raises the question of whether PVIs are necessary for the changes in PFC→MD neuron gamma synchrony we observed during the pre-decision period as mice learn the rule shift. To determine this, we next combined voltage recordings in PFC→MD neurons with optogenetic inhibition of PVI neurons (Fig. 4A-B). As in a previous study, we used red-shifted light for optogenetic inhibition so as to not interfere with TEMPO measurements^24^. We performed two days of rule-shifting to compare behavioral performance and synchrony measurements on Day 1, when no light was delivered for optogenetic inhibition, with Day 2, when we delivered 638nm light for optogenetic inhibition (Fig. 4C). We used experimental PV-Flp mice injected with AAV to drive Flp-dependent halorhodopsin (NpHR) expression in prefrontal PVIs (along with viruses for GEVI expression). We also used wild-type controls, which were Flp-negative and thus expressed GEVIs and fluorophores only (no NpHR). As expected, inhibiting prefrontal PVIs using 638nm light in NpHR+ PV-Flp mice impaired rule shift learning, as compared to NpHR-negative controls (Two-way ANOVA: Genotype: F_(1,15)_ = 5.12, p = 0.04; Stimulation Day: F_(1,15)_ = 1.22, p = 0.29; Fig. 4D). When we further analyzed performance separately for non-conflict and conflict trials, we found that PVI inhibition significantly increased perseveration, as evidenced by an increase in both correct choices on non-conflict trials and errors on conflict trials (Two-way ANOVA: Stimulation Day: F_(1,80)_ = 3.83, p = 0.05; Trial Type: F_(3,80)_ = 7.61, p < 0.001; Interaction: F_(3,80)_ = 6.326, p < 0.001; Fig. 4F). No increase in perseveration occurred on Day 2 in controls (Two-way ANOVA: Stimulation Day: F_(1,40)_ = 0.67, p = 0.42; Trial Type: F_(3,40)_ = 8.021, p < 0.001; Interaction: F_(3,40)_ = 0.41, p = 0.75; Fig. 4E).

**Figure 4.**
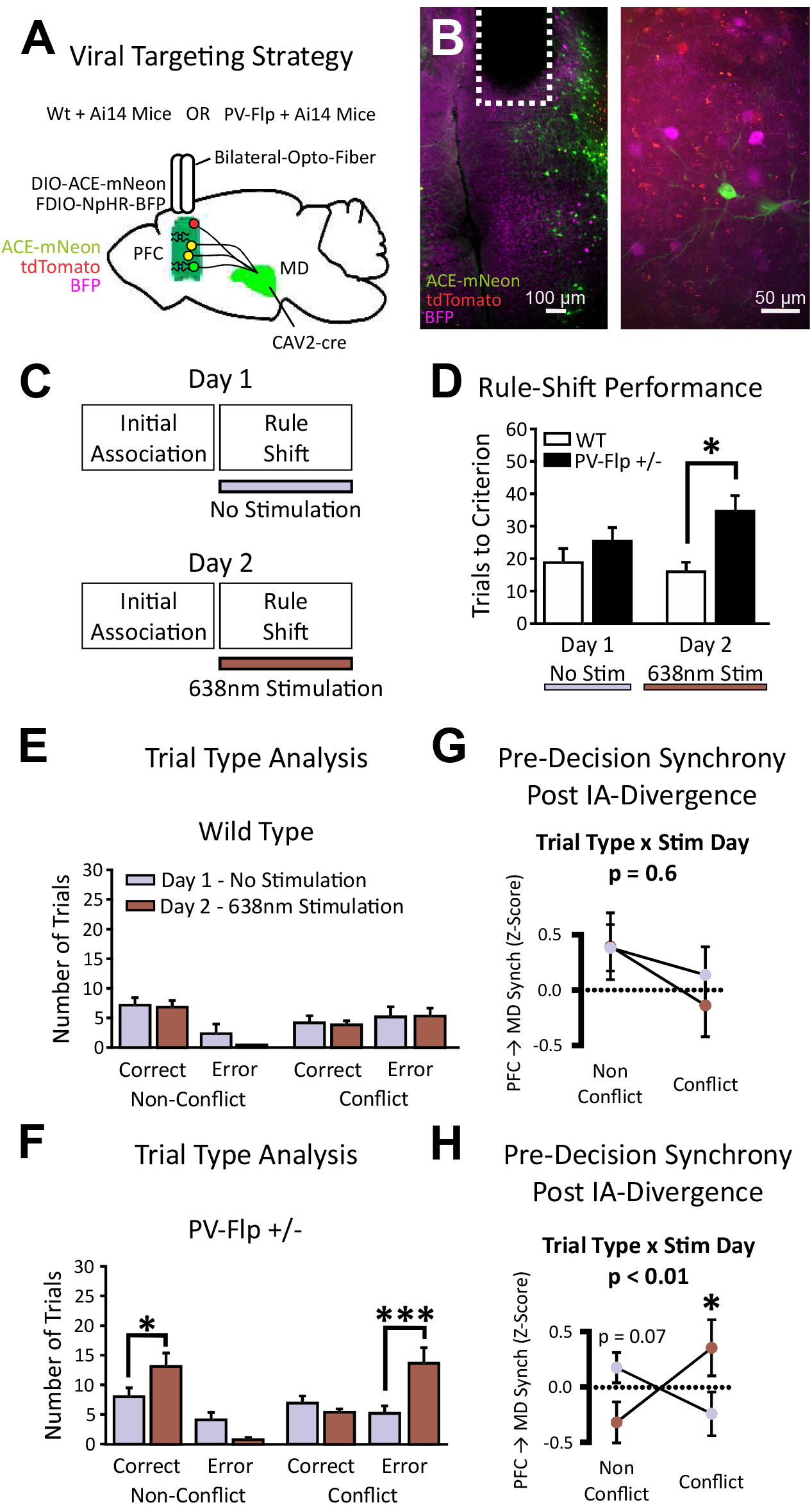
Optogenetic inhibition of prefrontal PV interneurons disrupts rule-shift learning and learning-induced patterns of synchrony in PFC→MD neurons. (A) Viral targeting strategy combining Cre-dependent expression of Ace-mNeon in retrogradely-labeled PFC→MD neurons with Flp-dependent expression of halorhodopsin (NpHR) in PV interneurons in the PFC of PV-Flp+/−; Ai14+/− mice. Bilateral fiber-optic implants allowed simultaneous voltage recording and optogenetic inhibition. (B) Representative histology. Ace-mNeon (green) and tdTomato (red) label Cre+ PFC→MD neurons. NpHR expression in PV interneurons was visualized by immunostaining for BFP with a 647 fluorophore (purple). The merged image shows combined colors in areas of signal overlap. (B) Experimental design: On Day 1, mice performed the initial association (IA) and rule shift (RS) phases of the task without light stimulation. On Day 2, 638nm light was delivered during the RS phase to inhibit PV interneurons. (C) Rule shift performance was significantly impaired in PV-Flp+ mice compared to wild-type controls during the stimulation session (*p<0.05). No significant differences were observed between groups on Day 1 (no stimulation). (D) Trial-type comparison in Flp-negative control mice showing the number of correct and error choices on conflict and non-conflict trials. No significant effects of stimulation were observed. (E) Trial-type comparison in PV-Flp+ mice showing that PV inhibition increased perseveration, with more correct responses on non-conflict trials and more errors on conflict trials (*p<0.05, ***p<0.001). (F) In Flp-negative controls, pre-decision interhemispheric gamma synchrony between PFC→MD neurons was unchanged across sessions. (G) In PV-Flp+ mice, optogenetic inhibition led to a significant interaction between stimulation day and trial type (p<0.01), with interhemispheric PFC→MD neuron gamma synchrony increasing on conflict trials (when it is normally low, *p<0.05) and trending toward a decrease on non-conflict trials (p=0.07).

Next, we examined PFC→MD synchrony during pre-decision periods following the IA-Divergent trial. We specifically quantified the interhemispheric synchrony between PFC→MD neurons in the left and right mPFC on both non-conflict and conflict trials depending on whether or not we delivered 638nm light for optogenetic inhibition. We did not see any significant effect of 638nm light in controls (Two-way ANOVA: Stimulation Day: F_(1,20)_ = 0.24, p = 0.63; Trial Type: F_(1,20)_ = 2.18, p = 0.16; Interaction: F_(1,20)_ = 0.30, p = 0.59; Fig. 4G). By contrast, in NpHR-expressing PV-Flp mice, we observed a significant interaction between day (Day 1 - no light vs. Day 2 - 638nm light for opto-inhibition) and trial type (Two-way ANOVA: Stimulation Day: F_(1,40)_ = 0.07, p = 0.80; Trial Type: F_(1,40)_ = 0.42, p = 0.52; Interaction: F_(1,40)_ = 7.61, p = 0.009; Fig. 4H). Opto-inhibition was specifically associated with a significant increase in interhemispheric PFC→MD neuron γ-synchrony on conflict trials (when synchrony is normally low; p = 0.035), and a trend towards decreased PFC→MD neuron γ-synchrony on non-conflict trials (when synchrony is normally high; p = 0.07). Thus, even though PVI-xH-γ-sync doesn’t increase during pre-decision periods following the IA-Divergent trials, PVIs still seem to mediate changes in PFC→MD γ-synchrony during these periods, as well as the associated transition in behavioral strategies.

### Changes in PFC→MD gamma synchrony occur because PVIs entrain ipsi- and contralateral PFC→MD neurons into behaviorally-specific phase configurations

If PVIs do not modulate their interhemispheric gamma synchrony during pre-decision periods, how do they modulate interhemispheric gamma synchrony in PFC→MD neurons? Furthermore, what exactly is the significance of interhemispheric gamma synchrony of PFC→MD neurons? One possibility that would address both of these questions is that during periods of interhemispheric PVI gamma synchrony, the ability of PVIs to entrain PFC→MD neurons into specific synchrony relationships might change. This could manifest as the sorts of changes in PFC→MD neuron interhemispheric synchrony we observed. To test this hypothesis, we sought to directly assess synchrony between PVIs and PFC→MD neurons. We did this by taking advantage of recent advances in TEMPO, namely methodologies that use spectrally-distinct GEVIs to simultaneously resolve voltage signals from two different cell types^30^, here PVIs and PFC→MD neurons. Specifically, we utilized three viruses to express Ace-mNeon2 specifically in prefrontal PVIs, the red-shifted GEVI Varnam2 specifically in PFC→MD neurons, and the reference fluorophore cyOFP nonspecifically across mPFC neurons (Fig. 5A-B). Because cyOFP is excited by blue light, but fluoresces in the amber range, this approach enabled us to separate cyOFP from Varnam fluorescence based on the fact that they are excited at two distinct modulation frequencies. Separated cyOFP signals can then be used to remove non-voltage signals for both GEVIs (Ace-mNeon and Varnam). Thus, this recently published approach^30^, enables the simultaneous measurement of 6 fluorescence channels (3 in each hemisphere) at high temporal resolutions in order to examine oscillatory dynamics between PVIs and PFC→MD neurons in the left and right prefrontal cortices (Fig. 5C).

**Figure 5.**
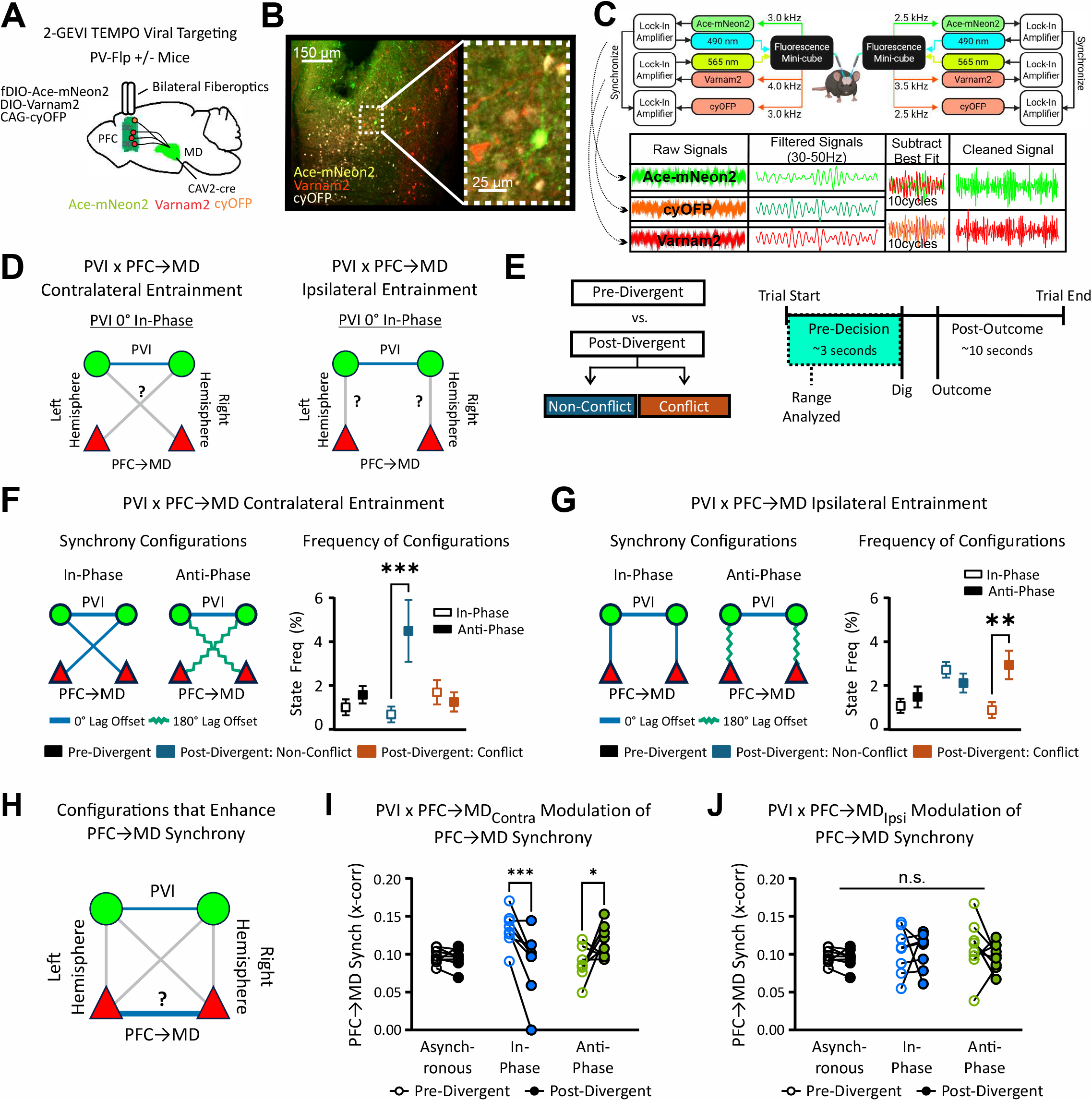
Dual-GEVI voltage measurements reveal that PVIs and PFC→MD neurons adopt distinct, behaviorally-specific synchrony configurations during learning. (A) Viral strategy for simultaneous GEVI recordings from PVIs and PFC→MD neurons. PV-Flp+/− mice were injected in mPFC with AAV2/PHP.eB-Ef1a-fDIO-Ace-mNeon2 (green) to label PV interneurons, AAVPHP.eB-Ef1a-DIO-Varnam2 (red) to label projection neurons, and AAV9-CAG-cyOFP (white) as a reference fluorophore. Retrograde CAV2-Cre injections in MD restricted Varnam expression to MD-projecting neurons. (B) Representative histology showing expression of Ace-mNeon in PV interneurons (green), Varnam2 in MD-projecting neurons (red), and cyOFP (reference fluorophore; white). (C) Hardware and analysis pipeline for extracting voltage signals. Dual LEDs (490 nm and 565 nm) were modulated by separate lock-in amplifiers to excite Ace-mNeon and Varnam-mRuby at separate frequencies; a third lock-in amplifier synchronized to the 490nm excitation frequency demodulated the red emission channel to separate the cyOFP reference signal. Raw fluorescence traces were filtered in the frequency band of interest, and we performed regression and subtraction using the reference cyOFP signal to remove shared non-voltage artifacts, and obtain cleaned Ace-mNeon and Varnam-mRuby voltage signals, which were then used for cross-correlation analyses. (D) Schematic illustrating possible synchrony configurations corresponding to combinations of phase relationships between PVIs and PFC→MD neurons. These were quantified by calculating cross-correlations between PVI signals and PFC→MD signals in the same and opposite hemispheres. (E) Schematic illustrating the specific behavioral epochs / trial types, and time window analyzed. The pre-decision period was analyzed on all trials, separated into three groups: 1) pre-divergent, 2) post-divergent / non-conflict, and 3) post-divergent / conflict trials. (F) Frequency of occurrence of specific synchrony configurations between PVIs and *contralateral* PFC→MD neurons. Post-divergent *non-conflict* trials showed a significant increase in the configuration characterized by anti-phase coupling between PVIs and contralateral PFC→MD neurons, compared to the configuration with in-phase coupling (**p<0.01). (G) Frequency of occurrence of synchrony configurations between PVIs and *ipsilateral* PFC→MD neurons. Post-divergent *conflict* trials were associated with an increase in anti-phase coupling, compared to in-phase coupling (**p<0.01). (H) Schematic illustrating our analysis, designed to determine the relationship between PVIs x PFC→MD neuron synchrony configurations and the level of interhemispheric PFC→MD neuron gamma synchrony. (I) Levels of interhemispheric PFC→MD neuron gamma synchrony observed when PVIs and *contralateral* PFC→MD neurons adopt specific synchrony configurations. After the IA-Divergent trial, interhemispheric PFC→MD neuron gamma synchrony increases for the anti-phase configuration but decreases for the in-phase configuration. (J) Levels of interhemispheric PFC→MD neuron gamma synchrony observed when PVIs and *ipsilateral* PFC→MD neurons adopt specific synchrony configurations. In this case, there are no significant differences in in interhemispheric PFC→MD neuron gamma synchrony before vs. after the IA-Divergent trial for any synchrony configuration.

We utilized this dual-GEVI approach to directly measure synchrony relationships between PVIs and PFC→MD neurons in either the same (PVI x PFC→MD_Ipsi_) or contralateral mPFC (PVI x PFC→MD_Contra_, Fig. 5D). Then we quantified and compared the occurrence of specific synchrony relationships during the pre-decision period prior to the IA-Divergent trial vs. the same period on non-conflict and conflict trials after the IA-Divergent trial (Fig. 5E). When PVI activity is in-phase across the hemispheres, the distribution of PVI x PFC→MD neuron phase relationships are largely bimodal, with peaks centered around 0 and 180 degrees. Therefore, we focused on configurations defined by either *in-phase* (near 0 degrees) or *anti-phase* (near 180 degrees) synchronization between these cell types. In other words, we first identified timepoints when PVIs exhibited in-phase interhemispheric synchrony, then further identified when this coincided with *in-phase* or *anti-phase* synchrony between PVIs and either *ipsilateral* or *contralateral* PFC→MD neurons.

Prior to the IA-Divergent trial (when mice do not differentiate behaviorally between conflict and non-conflict trials), all synchrony configurations occurred at similarly low levels. However, after the IA-Divergent trial, we observed a striking and behaviorally-specific enrichment of distinct synchrony configurations during the pre-decision period of conflict vs. non-conflict trials. Specifically, on *non-conflict* (but not on conflict) trials, there was a selective recruitment of *anti-phase* synchrony between PVIs and *contralateral* PFC→MD neurons (Two-way ANOVA: Synchrony Configuration: F_(1,42)_ = 5.33, p = 0.026; Trial Type: F_(2,42)_ = 2.03, p = 0.15; Interaction: F_(2,42)_ = 5.09, p = 0.011; Fig. 5F).

Conversely, on *conflict* (but not non-conflict) trials, there was a selective recruitment of *anti-phase* synchrony between PVIs and *ipsilateral* PFC→MD neurons (Two-way ANOVA: Synchrony Configuration: F_(1,42)_ = 2.88, p = 0.10; Trial Type: F_(2,42)_ = 3.26, p = 0.048; Interaction: F_(2,42)_ = 4.53, p = 0.017; Fig. 5G). This double dissociation shows that synchrony configurations change with learning, such that distinct patterns of PVI x PFC→MD neuron entrainment become selectively associated with specific trial types that involve different decision-making demands.

Finally, we asked whether the differential recruitment of specific synchrony configurations during the pre-decision period of non-conflict vs. conflict trials following the IA-Divergent trial could explain the differential modulation of interhemispheric PFC→MD neuron γ-synchrony we previously observed in Figure 3. For this, we quantified how each synchrony configuration affects interhemispheric PFC→MD neuron synchrony (Fig. 5H). We found that different PVI x PFC→MD_Contra_ configurations enhance PFC→MD synchrony depending on the stage of learning. *Prior to the IA-Divergent trial*, in-phase synchronization between PVIs and contralateral PFC→MD neurons was associated with enhanced PFC→MD interhemispheric gamma synchrony. However, *after the IA-Divergent trial*, anti-phase (but not in-phase) contralateral entrainment became associated with enhanced PFC→MD interhemispheric gamma synchrony (Two-way ANOVA: Synchrony Configuration: F_(2,42)_ = 1.56, p = 0.22; Trial Type: F_(1,42)_ = 0.70, p = 0.41; Interaction: F_(2,42)_ = 9.12, p < 0.001; Fig. 5I). By contrast PVI x PFC→MD_Ipsi_ configurations did not have any influence on PFC→MD synchrony (Two-way ANOVA: Synchrony Configuration: F_(2,42)_ = 0.52, p = 0.60; Trial Type: F_(1,42)_ = 0.53, p = 0.47; Interaction: F_(2,42)_ = 0.31, p = 0.73; Fig. 5J). This reveals that as mice diverge from the outdated IA strategy, prefrontal PVIs synchronize with each other and contralateral PFC→MD neurons populations during the pre-decision periods of non-conflict trials, leading to enhanced interhemispheric PFC→MD neuron gamma synchrony. In contrast, during the pre-decision period of conflict trials following the IA-Divergent trial, PVIs synchronize with each other and ipsilateral PFC→MD populations, but this fails to enhance interhemispheric PFC→MD neuron gamma synchrony. This explains why, as shown in Figure 3, interhemispheric PFC→MD neuron gamma synchrony diverges between non-conflict and conflict trials, and why, as shown in Figure 4, this relies on PVIs: this divergent pattern reflects the underlying synchrony configurations formed when PVIs in the two hemispheres synchronize with each other and entrain either ipsilateral (conflict trials) or contralateral (non-conflict trials) PFC→MD neurons.

## DISCUSSION

Motivated by prior findings from our laboratory that interhemispheric gamma synchrony between prefrontal PVIs (PVI-xH-γ-sync) increases after mice make rule shift errors, here we sought to understand the function of these increases, by examining whether they are tied to specific aspects of cognitive flexibility and transmitted to other cell types. We find that PVI-xH-γ-sync increases specifically during the post-outcome period of trials associated with transitions in behavioral strategies. We do not find that increases in PVI-xH-γ-sync during the post-outcome periods propagate to projection neurons. However, when mice ultimately stop following the original rule, gamma synchrony during the pre-decision period changes for mPFC neurons which project to the MD thalamus, but not those projecting to dorsal striatum. Moreover, these changes distinguish between conflict and non-conflict trials: a distinction which is initially behaviorally-irrelevant, but becomes relevant during the rule shift portion of the task. Thus, changes in gamma synchrony seem to be a neural signal for evolving behavioral contingencies. Specifically, we find that on non-conflict trials, periods of PV-xH-γ-sync are associated with increased entrainment between PVIs and *contralateral* PFC→MD neurons, whereas on conflict trials, periods of PV-xH-γ-sync are associated with increased PVI entrainment with *ipsilateral* PFC→MD neurons. These results show for the first time that prefrontal gamma oscillations comprise multiple distinct local and long-range synchrony configurations which each involve specific cell types and subserve different, dissociable aspects of flexible behavior.

### New ways of thinking about gamma oscillations

The classic notion of a gamma oscillation is that rhythmic activity in inhibitory interneurons entrains local excitatory neurons, synchronizing activity throughout a circuit. This was demonstrated most clearly in seminal experiments which induced ∼40Hz oscillations in hippocampal and neocortical slices^31^. This concept is the foundation for many hypothesized functions of gamma oscillations, most notably communication through coherence, which posits that during gamma oscillations, rhythmic output from inhibitory neurons creates narrow windows of excitability in local excitatory neurons, such that the proper alignment (coherence) of these windows between regions is necessary for them to interact effectively^27,32^. In contrast to this classic notion, here we observe gamma-frequency synchronization between PVIs in the left and right mPFC without necessarily observing similar synchronization in two different populations of projection neurons. When we do observe increases in interhemispheric gamma-frequency synchronization in MD-projecting neurons, this does not generalize to another class of projection neurons. Even more surprisingly, when PVIs synchronize across hemispheres, they can entrain MD-projecting neurons, but on non-conflict trials they mainly entrain the MD-projecting neurons in the contralateral, but not ipsilateral, mPFC. Together, these results demonstrate that synchronized gamma-frequency PVI activity can co-exist with multiple possible patterns of synchronization which entrain specific subsets of local or distant projection neuron populations.

An additional note is that when specific patterns of entrainment between PVIs and projection neurons become enriched, this tends to involve *antiphase* synchronization between these two cell types, rather than the short phase delays that would be predicted by pyramidal neuron-interneuron network gamma (‘PING’) models in which pyramidal neuron firing rapidly recruits inhibitory neuron firing^27,33^. That said, GEVI kinetics can introduce temporal shifts in rhythmic activity, and under some conditions these can depend on the combination of indicator and cell type^30^. Thus, the precise magnitude of the phase difference between these two populations could be shifted relative to what we calculated using GEVIs.

Recently there have been concerns about using gamma oscillations as readouts of cognitive processes since it is becoming increasingly clear that measuring gamma oscillations at the LFP level amalgamates multiple circuit processes, each involving unique sets of inputs, and may thus obfuscate these individual underlying processes^12^. Our results support the interpretation that gamma oscillations must be resolved in a circuit-specific rather than bulk manner, and further illustrate how this can be done through the use of GEVI-based methods such as one and two-color TEMPO^26,30^. Our observations that these methods reveal increases in γ-synchrony which manifest in PFC→MD neurons, but not in neighboring neurons projecting to dorsal striatum, or which occur selectively between PVIs and either ipsilateral or contralateral PFC→MD neurons depending on levels of cognitive conflict, further validate that these findings reflect cell type-specific voltage signals, not non-specific artifacts related to fiber bending or blood oxygenation. Notably, whereas we used a method that we had developed when this study began to minimize the influence of artifacts on TEMPO signals, more recent methods for improving optical sensitivity and the computational unmixing of artifactual and neural voltage signals^30^ might resolve gamma-frequency phenomena in even greater detail and/or with even greater detection fidelity.

### Different synchrony configurations subserve distinct functions

We find that PVI-xH-γ-sync increases during the post-outcome period on trials that correspond to behavioral transitions: the IA-Divergent and RS-Acquisition trials. This pattern of synchronization is not necessarily associated with changes in gamma synchrony in neurons projecting to MD thalamus or dorsal striatum. This suggests that gamma synchrony involving PVIs may contribute to learning-related processes, possibly by regulating synaptic plasticity at PVI synapses^35^ or elsewhere in the microcircuit. By contrast, during the pre-decision period, we find that overall levels of PVI-xH-γ-sync do not change, but that periods of PVI-xH-γ-sync coincide with ipsilateral PVI x PFC→MD neuron γ-synchrony on conflict trials, and with contralateral PVI x PFC→MD neuron γ-synchrony on non-conflict trials. Conflict trials are associated with the greatest requirement for cognitive control, specifically the need to suppress perseverative responses in order to choose the bowl which contains the cue that signals reward during the RS. Thus, antiphase γ-synchronization between PVIs and ipsilateral PFC→MD neuron synchrony may serve to disrupt outdated activity patterns and/or facilitate the emergence of new activity patterns within PFC-MD loops which have been implicated in flexible behavior^5,36,37^. It is notable that non-conflict trials recruit a unique pattern of synchronization, which suggests that prefrontal circuits engage distinct mechanisms on these trials compared to trials on which they follow the initial association strategy, even though their behavioral output (choice behavior) is ultimately the same. The function of PVI x *contralateral* PFC→MD neuron synchronization on these trials is not entirely clear but might serve to prevent choices which are consistent with the initial association strategy from inappropriately reinforcing or reinstating that strategy. Altogether these findings indicate that in this task gamma synchrony does not perform just one function (e.g., promoting learning), but rather that different synchrony configurations (all of which include in-phase interhemispheric PVI γ-synchronization) are dynamically recruited to subserve specific behavioral needs.

Identifying these specific functions for gamma synchrony is important for many reasons. First, the idea that gamma oscillations contribute meaningfully to brain function has long been a major controversy in systems neuroscience^38,39^. Our previous work has established that gamma synchrony is necessary for rule shift learning^22^ but linking it more specifically to particular aspects of this task (learning new representations, encoding behavioral contingencies, shaping choice behavior) is a key next step in elucidating the specific information mechanisms through which gamma oscillations act.

Along these lines, our finding that learning involves increases in PVI x contralateral PFC→MD neuron γ-synchrony on non-conflict trials is particularly intriguing, as we have previously shown that callosal PV projections mainly target PFC→MD neurons and that stimulating or inhibiting these projections can lead to long-lasting increases or decreases in cognitive flexibility, respectively. Taken together these findings are consistent with a mechanism in which callosal PV synapses onto PFC→MD undergo potentiation during rule shift learning, possibly triggered by PVI-xH-γ-sync during the post-outcome period of behavioral transitions. Indeed, recent work from our lab has found evidence for this type of synaptic potentiation during rule shift learning^40^.

### Clinical implications

Identifying specific functions of gamma synchrony in this task may also inform the development of novel therapeutics for cognitive deficits in schizophrenia and related conditions. Enhancing cognitive function in psychiatric disease is a paramount goal of mental health research but severely limited by our current understanding of the mechanisms underlying normal cognition. Enhancing task-relevant gamma oscillations in the prefrontal cortex has strong therapeutic potential^9–11^, however it has become increasingly clear that gamma oscillations, unlike slower neural rhythms, exhibit complex dynamics which need to be understood before effective therapies can be developed^12,41^. For example, simply measuring the overall power of gamma oscillations in a non-specific manner is unlikely to be useful as a translational biomarker or for validating or optimizing novel therapeutics. Our results suggest that to be meaningful, such measurements need to account for cell types, phase relationships, and behavioral contingencies.

### Behavioral contingencies regulate gamma synchrony in PFC→MD neurons

In addition to having temporal and circuit specificity, we also discovered that gamma synchrony manifests in a behavioral contingency-specific manner – even during the same period of learning (after the IA-Divergent trial) and behavioral epoch (pre-decision period). Specifically, after mice diverge from their old strategy, PVIs and PFC→MD neurons exhibit divergent patterns of synchronization on conflict vs. non-conflict trials. This aligns with other studies suggesting that PVI-mediated inhibition is necessary for normal modulation of prefrontal gamma activity during the pre-decision period^42^, and that PFC→MD communication plays an important role in guiding choice selection^5,29^. Interestingly, one of these prior studies^5^ found that periods of behavioral uncertainty can recruit two distinct MD circuits which differentially influence the PFC. When uncertainty is related to sparse information, dopamine D2 receptor-expressing MD neurons can excite PFC VIP interneurons to amplify PFC activity. By contrast during periods of dense, but conflicting, information, GRIK4-expressing MD neurons excite PFC PVIs to suppress PFC activity. Differential activity of these MD→PFC pathways may underlie the divergent PVI x PFC→MD neurons synchrony dynamics we observed on conflict vs. non-conflict trials.

## Conclusion

Using voltage indicators and optogenetics, we find that PVIs dynamically engage multiple distinct synchrony configurations, each of which involves specific phase relationships between local and distributed cell types and emerges in a behaviorally-specific manner. These findings upend conventional notions of gamma oscillations as singular, microcircuit-wide phenomena, revealing new dimensions and illustrating how GEVI-based approaches can resolve multiple functionally-distinct synchrony configurations occurring within the same circuit. Future experiments should explore relationships between these synchrony configurations and additional cell types, synaptic plasticity mechanisms, information processing in PFC-MD thalamocortical loops, pathological states related to schizophrenia, and novel therapeutic interventions.

## ACKNOWLEDGEMENTS

This work was supported by BBRF Young Investigator Award #31117 (to A.P.), and NIH grants R01NS116594 (to V.S.S.), R01MH117961 (to V.S.S., M.J.S. and M. Kheirbek), R01MH106507 (to V.S.S.), R01NS124590 (to M.J.S. and I. Soltesz), and U01NS120822 (to M.J.S. and G. Vasan).

## DECLARATION OF INTERESTS

M.J.S. is co-author of a U.S. patent, and S.H. and M.J.S. have pending patent applications related to the work in this paper.

## METHODS

### Animals and Housing Conditions

Animals of both male and female sexes were used for all experiments. Mice were normally group-housed in a 12hr light / dark cycle with ad libitum access to food. For rule shift behaviors animals were moved to a reverse light / dark schedule and singly housed for the duration of behavior. During this time animals were food restricted (as described below) over the course of ∼ 1 week to 85% of baseline weight.

For experiments examining GEVI expression in PVI (Fig 2), animals of PV-Cre (Jax; 017320) line were crossed with Ai14 (Jax; 007914) to generate PV-Cre^+/-^/ Ai14^+/-^mice enabling expression of Ace-mNeon and TdTomato in prefrontal PVI. For experiments examining GEVI expression in PFC-MD or PFC-DS neurons (Fig 3), Ai14 mice were used for all experiments. For experiments combining GEVI expression in PFC-MD neurons with fDIO-NpHR3.0-BFP expression in prefrontal PVI, animals of the PV-Flp (Jax; 022730) line were crossed with Ai14 to generate PV-Flp^+/-^/ Ai14^+/-^mice enabling expression of Ace-mNeon and TdTomato in PFC-MD neurons and NpHR3.0-BFP in prefrontal PVI (Fig 4). PV-Flp^-/-^/ Ai14^+/-^from the same litters were used as wild-type controls. For experiments using the dual-color GEVIs Ace-mNeon2 and Varnam2, PV-Flp^+/-^(Jax; 022730) mice were used.

### Viral vectors

#### Optogenetic Experiments

We first obtained the AAV plasmid pAAV-nEF-Coff/Fon-NpHR3.3-EYFP from addgene (Addgene plasmid # 137154) and had Virovek replace the EYFP genetic sequence for BFP. protein sequence. generate AAV8-nEF-Coff/Fon-NpHR3.3-BFP

#### Single-GEVI TEMPO Experiments

We obtained the AAV1-CAG-DIO-Ace2N-4AA-mNeon^26,43^ plasmid from the Schnitzer lab and manufactured the virus through Virovek at a working titer of 2.8E13 GC/ml. We have used this virus extensively in previous studies^22,24,34^.

#### Dual-GEVI TEMPO Experiments

The Schnitzer lab provided the AAV2/PHP.eB-Ef1a-DIO-Varnam2-WPRE (plasmid #240055)^30^. We obtained the AAV plasmid pAAV-Ef1a-fDIO-Varnam2-WPRE by cloning the genetic sequence encoding the fluorescent GEVI proteins Varnam2 into the AAV backbone pAAV-EF1a-fDIO-Cre (Addgene plasmid # 121675) by inserting the Varnam2 protein sequences in place of the Cre sequence between NheI and AscI targeting sequences.

All sequences for red and green FRET-opsin GEVIs included the endoplasmic reticulum (ER) export sequence and Golgi export trafficking signal (TS). We confirmed that vectors had accurate sequences using Sanger sequencing (Quintara Biosciences). We used the QIAGEN Plasmid Plus Maxi Kit (#12963, QIAGEN) to amplify all plasmids prior to viral packaging. HHMI/Janelia Viral Tools manufactured all viruses used in the dual-GEVI experiments.

We expressed green GEVIs using AAV2/PHP.eB-Ef1a-fDIO-Ace-mNeon2-WPRE (3.6E13 GC/mL).

We expressed red GEVIs using AAV2/PHP.eB-Ef1a-DIO-Varnam2-WPRE (1.18E13 GC/ml).

We expressed reference fluor via AAV2/PHP.eB-CAG-cyOFP (5.6E13 GC/mL) (Addgene plasmid # 240050).

### Rule Shift Behavior

See figure 1D-E. Methods are adapted from previous work^22^. In short, mice are first food restricted to 85% of baseline weight. Mice then undergo habituation where they are trained to dig in one of two bowls containing complex cues, each consisting of an odor and texture modality. The bowl pairs are organized into 2 presentations: 1. Litter + Coriander and Sand + Garlic (Pair 1) or 2. Litter + Galic and Sand + Coriander (Pair 2). During habituation cues are evenly rewarded and no cue-reward association is formed. Mice learn to do this relatively fast, and habituation is completed over the course of 10 presentations (trials). On the next day mice perform the task which is broken into two parts: Initial Association (IA) – mice are trained to attend a specific modality and discriminate between the two cues to receive a reward (e.g. IA Rule: Sand: mice must dig in either Sand + Garlic or Sand + Coriander bowls depending on presentation). If mice dig in the correct bowl, a reward (peanut butter chip) is delivered to the mouse using forceps and then they are placed in a holding cage for ∼30 seconds while the next trial is prepared. If mice dig in the incorrect bowl, then the experimenter removes the correct bowl (to signal an error) and mice are placed in a holding cage for ∼90 seconds. After animals achieve a behavioral criterion of 8/10 correct choices in a row, animals directly begin the next portion: Rule Shift. During the Rule Shift, the reward-cue pairing is shifted across modalities (e.g. IA Rule: Sand -> RS Rule: Garlic: mice must dig in either Sand + Garlic or Litter + Garlic). This shift is uncued and requires animals to ignore the previous rule’s modality while attending the new modality which is considered an extradimensional shift. Note that 50% of the time an animal can use the IA rule and still get a correct choice (e.g. IA Rule: Sand -> RS Rule: Garlic, Sand + Garlic choice), these trials are considered ‘Non-Conflict’ trials, whereas the other trials (e.g. requiring Litter + Garlic choice) are considered ‘Conflict’ trials since animals must actively ignore the IA rule. Correct and error trials are handled the same as in IA and animals continue the behavior until they once more reach a criterion of 8/10 trials.

### Trial Classification Algorithm and Behavioral Scoring

All behavioral videos were captured through the Tucker-Davis Technology’s data acquisition software, Synapse, which ensures frames are appropriately time-locked to neural data. All behavioral events (trial start, trial end, dig, outcome) were scored using a custom MATLAB script.

We categorize behavior on each trial as belonging to one of three strategy categories: **stable, exploratory**, or **divergent**. Stable strategy blocks follow one consistent choice policy (e.g. reliably choosing a specific cue or side), whereas exploratory blocks do not.

Divergent trials mark the transitions between stable strategies or from stable to exploratory behavior.

We first assume that mice follow one of six basic policies – four cue-based (e.g. litter, sand, garlic, coriander) or side-based (left or right). Each trial was scored using three metrics:

1. Past policy stability (PPS): The fraction of the six preceding choices that employ a given policy.
2. Future policy stability (FPS): Fraction of the current and five subsequent choices that employ a given policy.
3. Current-trial predictability (CTP): The probability of the current choice based on recent choice history.

For PPS and FPS, we weighed the three most recent (or subsequent) trials by an extra 50% to emphasize local stability. To calculate CTP, we fit a logistic regression model and used it to calculate the probability of the current trial choice given the six previous trial choices.

Specifically, we utilized a logistic regression model to predict the bowl chosen on the current trial, with binary input variables for whether the cue/side chosen on previous trials matches those in each bowl on the current trial. We fit the model on all trials pooled across animals using 10-fold cross-validation and ElasticNet regularization. We define CTP as the model’s conversion probability of the actual choice (Supp. Fig. 1A-B).

We perform K-nearest neighbors (KNN) clustering on these three metrics (PPS, FPS, CTP; Supp. Fig. 1C-E) and use the resulting cluster boundaries to set thresholds for inclusion in each strategy class. Classification thresholds were as follows:

- Stable trials: PPS > 0.9 and either CTP > 0.4 or FPS > 0.9
- Divergent trials: PPS > 0.9 and CTP ≤ 0.4
- Exploratory trials: Low PPS, FPS, and CTP

To evaluate our classification strategy, we simulated trial choices from randomly generated fixed-length blocks of stable (cue- or side-based) and exploratory (randomized) behavior. We varied the simulated strategy length (number of trials per block) and window size (number of trials used to compute PPS, FPS, and CTP). 50,000 trials were generated per condition. We assessed performance using four metrics: switch detection accuracy (correct classification of transitions), sensitivity (proportion of true switches detected), precision (proportion of predicted switches that were correct) (Supp. Fig. 1F–H), and classification accuracy (fraction of trials correctly labeled by the algorithm) (Supp. Fig. 1I).

Classification performance improved with longer strategy blocks and moderate window sizes. Precision was lower for shorter blocks where behavior was more erratic, occasionally leading to over-identification of transitions. Accuracy, sensitivity, and overall classification remained high across most conditions. A window size of 6 (used in the final analysis) provided a strong balance across all metrics. This choice is further supported by the task structure: animals were required to complete at least 8 correct trials out of 10 during both the Initial Association and Rule Shift phases, imposing stable behavior. Moreover, our analysis focused on IA-Divergent and RS-Acquisition trials—epochs tied to these criteria— effectively selecting for longer blocks. A 6-trial window thus offers a principled and conservative estimate for capturing transitions in cognitive strategy. Two examples of the classification strategy applied to a full session of behavior are provided (Supp. Fig 1J-K).

### Stereotactic Surgeries

Male and female mice were anaesthetized using isoflurane (2.5% induction, 1.2–1.5% maintenance, in 95% oxygen) and placed in a stereotaxic frame (David Kopf Instruments). Body temperature was maintained using a heating pad. An incision was made to expose the skull for stereotaxic alignment using bregma and lambda as vertical references. The scalp and periosteum were removed from the dorsal surface of the skull and scored with a scalpel to improve implant adhesion. Viruses were infused at 100–150 nl/min through a 35-gauge, beveled injection needle (World Precision Instruments) using a microsyringe pump (World Precision Instruments, UMP3 UltraMicroPump). After infusion, the needle was kept at the injection site for 5–10 min and then slowly withdrawn. After surgery, mice were allowed to recover until ambulatory on a heated pad, then returned to their home cage. For all experiments, we waited at least 5 weeks to allow for virus expression prior to beginning behavioral experiments.

For experiments investigating γ-Synchrony in prefrontal PVI (Fig 2), PV-Cre^+/-^/ Ai14^+/-^mice were injected bilaterally into the mPFC (+1.7 AP, +/-0.3ML, three depths: −2.5, −2.25, −2 DV) with 200nL (per depth) of AAV1-CAG-DIO-Ace2N-4AA-mNeon (Virovek). After injection of virus, two mono fiber-optic cannulas (Doric Lenses; MFC_400/430-0.48_2.8mm_ZF1.25_FLT) were lowered into the left and right prefrontal cortices (+1.7 AP, +/-0.76ML, −2.13DV) at a 12-degree angle.

For experiments investigating γ-Synchrony in deep layer PFC neurons targeting either the MD or DS (Figs 2 and 3), Ai14^+/-^mice were injected bilaterally into the mPFC (+1.7 AP, +/-0.5ML, three depths: −2.5, −2.25, −2 DV) with 200nL (per depth) of AAV1-CAG-DIO-Ace2N-4AA-mNeon (Virovek). In addition, mice were injected bilaterally into either the mediodorsal thalamus (−1.7 AP, +/-0.35ML, −3.5DV) or dorsal striatum (+0.9 AP, +/-1.0ML, - 3.0 DV) with 350nL (per site) with CAV2-Cre (Plateforme de Vectorologie de Montpellier). After injection of virus, two mono fiber-optic cannulas (Doric Lenses; MFC_400/430-0.48_2.8mm_ZF1.25_FLT) were lowered into the left and right prefrontal cortices (+1.7 AP, +/-0.8ML, −2.2DV) at a 12-degree angle.

For experiments combining the investigation of γ-Synchrony in MD-projecting neurons with inhibition of prefrontal PVI (Fig 4), PV-Flp^+/-^/ Ai14^+/-^mice were injected bilaterally into the mPFC (+1.7 AP, +/-0.5ML, three depths: −2.5, −2.25, −2 DV) with 250nL (per step) of a 1:1 ratio of AAV1-CAG-DIO-Ace2N-4AA-mNeon (Virovek) and AAV8-nEF-Coff/Fon-NpHR3.3-BFP (Virovek). In addition, mice were injected bilaterally into the mediodorsal thalamus (−1.7 AP, +/-0.35ML, −3.5DV) with 350nL (per site) with CAV2-Cre (Plateforme de Vectorologie de Montpellier). After injection of virus, two mono fiber-optic cannulas (Doric Lenses; MFC_400/430-0.48_2.8mm_ZF1.25_FLT) were lowered into the left and right prefrontal cortices (+1.7 AP, +/-0.8ML, −2.2DV) at a 12-degree angle.

For experiments measuring γ-Synchrony in PVI and PDC→MD neurons simultaneously (Fig 5), PV-Flp^+/-^mice were injected bilaterally into the mPFC (+1.7 AP, +/-0.5ML, three depths: −2.5, −2.25, −2 DV) with 250nL (per depth) of a combination of AAV1-CAG-DIO-Ace2N-4AA-mNeon (Virovek), Ef1a-fDio-VA2.034-WPRE (Schnitzer Lab), and CAG-cyOFP-H-WPRE (Schnitzer Lab). In addition, mice were injected bilaterally into the mediodorsal thalamus (−1.7 AP, +/-0.35ML, −3.5DV) with 350nL (per site) with CAV2-Cre (Plateforme de Vectorologie de Montpellier). After injection of virus, two mono fiber-optic cannulas (Doric Lenses; MFC_400/430-0.48_2.8mm_ZF1.25_FLT) were lowered into the left and right prefrontal cortices (+1.7 AP, +/-0.8ML, −2.2DV) at a 12-degree angle.

### Instrumentation for single and dual cell-types fiber-optic TEMPO

Methods for single-cell type TEMPO recording (Fig 2) are described in detail in previous work^24^ but are described in brief here. Per recording site, activation of Ace-mNeon and tdTomato fluorophore was accomplished using two fiber-coupled LEDS (Thorlabs; M490Fr and M565F3) controlled by an LED Driver (Thorlabs; DC4104) and each LED was modulated by a sinusoidal signal provided by a lock-in amplifier (Stanford Research Systems; SR860) at a specific frequency to allow for demodulation of each signal. Both LEDs were fed into a ‘mini-cube’ (Doric Lenses, FMC5_E1(460-490)_F1(500-540)_E2(555-570)_F2(580-600)_S) and then transmitted into the animal through a mono fiber-optic patch cord (Doric Lenses, MFP_400/430/1100-0.48_2m_FC-ZF1.25) attached to the mono fiber-optic cannula implanted on the animal during surgery. GEVI emissions were captured through the same ‘mini-cube’ and fed into a photoreceiver (FEMTO OE-200-SI-FC) which sends the resulting signal back to the lock-in amplifier. All four signals are then demodulated and then digitized by a multichannel real-time signal processor (Tucker-Davis Technologies, RX-8). The commercial software Synapse (Tucker-Davis Technologies) was used for data acquisition.

For experiments that combined GEVI recordings with optogenetic inhibition of PVI, the system was the same as described above, however a different ‘mini-cube’ was used that had an additional opto-port (Doric Lenses, FMC6_E1(460-490)_F1(500-540)_E2(555-570)_F2(580-600)_O(628-642)_S) which was connected to a 638nm laser (Doric Lenses). Laser power was set to 2.5mW and measured using a ThorLabs light meter (PM100D) at the fiber tip.

For experiments utilizing the dual-cell type fiber optic TEMPO measurements from two brain areas concurrently, we used the methods described in Haziza et al^30^. In brief, we targeted two neuronal populations using the two GEVIs, Ace-mNeon2 and Varnam2^44^, and replaced the tdTomato reference fluor by the long-stock shift fluorescent proteins cyOFP^45^. We utilized the same system described above, however added two additional lock-in amplifiers (Stanford Research Systems; SR860). Each additional lock-in amplifier’s frequency was set to external modulation and received input from the two respective lock-ins modulating the green LED. However, these two additional lock-in amplifiers received input from the ‘red channel’ photoreceiver (mixture of Varnam2 and cyOFP signals).

Effectively, this allowed these lock-ins to pick up activity of the cyOFP reference fluorophore, which is activated by the green LED, but emits at 590nm. cyOFP was then utilized as the reference fluorophore to remove non-voltage signals from both signals arising from Ace-mNeon2 and Varnam2. All six signals are then demodulated and digitized as above.

#### Analysis of fiber-optic TEMPO data

Data from all four channels were acquired using Tucker-Davis Technologies Synapse software and converted into a MATLAB structure using the SEV2Mat utility (Tucker-Davis Technologies). Signals were first band-pass filtered within the frequency range of interest (30–50 Hz for gamma). Signal cleaning was performed by applying linear regression within 250 ms bins between the Ace-mNeon and tdTomato signals from each hemisphere and subtracting the shared noise component from the Ace-mNeon signal, resulting in a voltage signal corrected for shared, non-voltage-related activity.

To calculate cross-hemispheric gamma synchrony, we computed the zero-lag Pearson correlation coefficient between the cleaned voltage signals recorded from the left and right prefrontal hemispheres in 250 ms bins across the full recording session. Additionally, to examine phase relationships while avoiding the high computational cost of traditional cross-correlation functions, we linearly shifted one signal relative to the other across a single 40 Hz gamma cycle (–180° to +180°) in 22.5° increments. For each shift, correlations were computed within the same 250 ms bins. This approach allowed us to categorize synchrony as **In-Phase** (when the maximal Pearson correlation coefficient fell between – 45° and +45°) or **Anti-Phase** (when the maximal correlation fell between +135° and +225°). In-Phase synchrony was used for further analyses unless otherwise specified. For z-scoring, a baseline period of 10 minutes prior to the start of behavior was used to normalize the data.

For experiments utilizing dual-color GEVIs, synchrony between the two populations were calculated in a similar manner, however six channels were extracted (Left Hemisphere: Ace-mNeon2, Varnam2, cyOFP; Right Hemisphere: Ace-mNeon2, Varnam2, cyOFP) and signal cleaning utilized CAG-cyOFP as the reference fluorophore and was used to ‘clean’ both voltage signals. γ-Synchrony was then calculated in the same way as described above.

To determine how synchrony configurations related to cross-hemispheric γ-synchrony in PFC→MD neurons (Fig. 5H–J), we identified all timepoints at which each specific synchrony configuration occurred (as defined in Fig. 5F–G). For these timepoints, we then calculated In-Phase γ-synchrony in PFC→MD neurons in the same manner described above, separately for Pre- and Post-Divergent trials.

#### Validation of Analysis Pipelines used to Calculate Gamma Synchrony

To validate our ability to identify periods of gamma synchrony following our signal filtering, cleaning, and processing steps we used simulated data with known periods of gamma synchrony embedded with realistic noise. Simulated signals were composed of gamma oscillations combined with shared, independent, and broadband noise. The Ace-mNeon channels contained both signal and noise, while tdTomato channels contained only noise components. We generated three independent sets of data (“subjects”) which included a 15-minute baseline and 15 minutes of ‘task data’ broken into 5 trials per set. In each set, two trials contained periods of strong in-phase gamma synchrony specifically during the post-outcome period, while all other epochs contained asynchronous phase relationships. Applying our full analysis pipeline to this dataset successfully recovered elevated gamma synchrony during the defined post-outcome windows of the synchronized trials (Supp. Fig. 2A-B).

To validate that our analysis pipeline does not produce false positives, we calculated gamma synchrony in our real data (PVI dataset) in which the phase relationship between the left and right Ace-mNeon signals was abolished. This was achieved by applying a Fourier-based phase-scrambling method^46^ to one of the signals, preserving its power spectrum while randomizing phase. Under these conditions, we found that the increase in PVI-Synch observed on IA-Divergent and RS-Acquisition trials (Fig. 2E) was abolished (Supp. Fig. 2D), thus demonstrating that our synchrony measure specifically captures phase-related increases in gamma-synchrony.

Finally, to confirm that the observed synchrony was attributable to voltage-dependent activity rather than non-voltage-related noise, we calculated gamma synchrony between the left hemisphere Ace-mNeon signal and the right hemisphere tdTomato signal. As expected, we observed no significant synchrony between these signals, reinforcing that the increased synchrony reported in Fig. 2 is specifically driven by correlations in voltage dynamics (see Supp. Fig. 2E–F).

### Histological Tissue Validation

Following experiments animals were anesthetized with an i.p. injection of Euthasol and transcardially perfused with 30 ml of ice-cold 0.01 M PBS followed by 30 ml of ice-cold 4% paraformaldehyde in PBS. Brains were extracted and stored in 4% paraformaldehyde for 24 h at 4 °C before being stored in PBS. Slices 70–100 μm thick were obtained on a Leica VT100S and mounted on slides. All imaging was performed on an Olympus MVX10,Keyence BZ-X All-in-One Fluorescence Microscope, or Nikon Crest LFOV Spinning Disk/ C2 Confocal. All mice were verified to have virus-driven expression and optical fibers located in the mPFC.

---

**Supplementary Figure 1.**
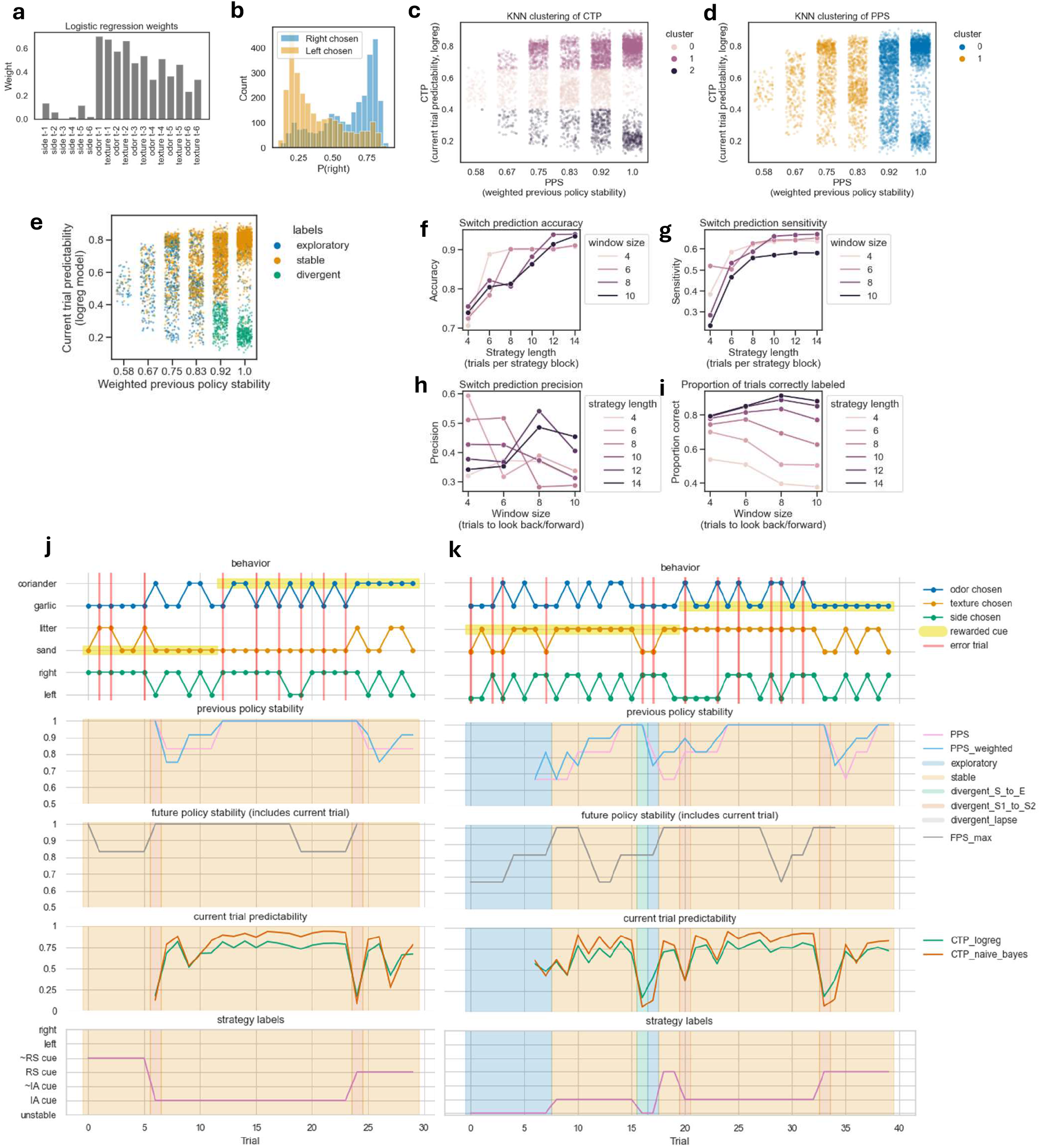
Details of behavioral strategy classification and validation. Logistic regression outputs and simulation-based validation of behavioral strategy classification, illustrating how CTP, PPS, and clustering contribute to trial classification and how model performance varies across strategy lengths and window sizes. A) Coefficients from the logistic regression model used to calculate current-trial predictability (CTP) based on previous trial choices. B) Histogram of conversion probabilities for choosing the right bowl, as predicted by the logistic regression model, colored by the actual choice made on the current trial. C) Scatterplot of past-policy stability (PPS) versus CTP, with points colored according to k-nearest neighbor (kNN) clustering based on CTP. Data shown as individual trials pooled across animals. D) Same as (C), but colored according to clustering based on PPS. E) Same as (C), but colored according to final classification labels (stable, exploratory, or divergent). F) Classification accuracy for detecting strategy switches on simulated data as a function of block length, colored by window size used for classification. G) Same as (F), but showing sensitivity (true positive rate) for switch detection. H) Same as (F), but showing prediction precision (proportion of predicted switches that were correct) as a function of window size, colored by strategy length. I) Proportion of trials correctly labeled by the classifier, shown as a function of window size and colored by strategy length. J–K) Two example behavior sessions. Top panels show the cue and side chosen on each trial. Bottom panels show PPS, FPS, CTP, and the inferred policy, with overlays indicating trial-by-trial classification labels (stable, exploratory, divergent) according to the model.

**Supplementary Figure 2.**
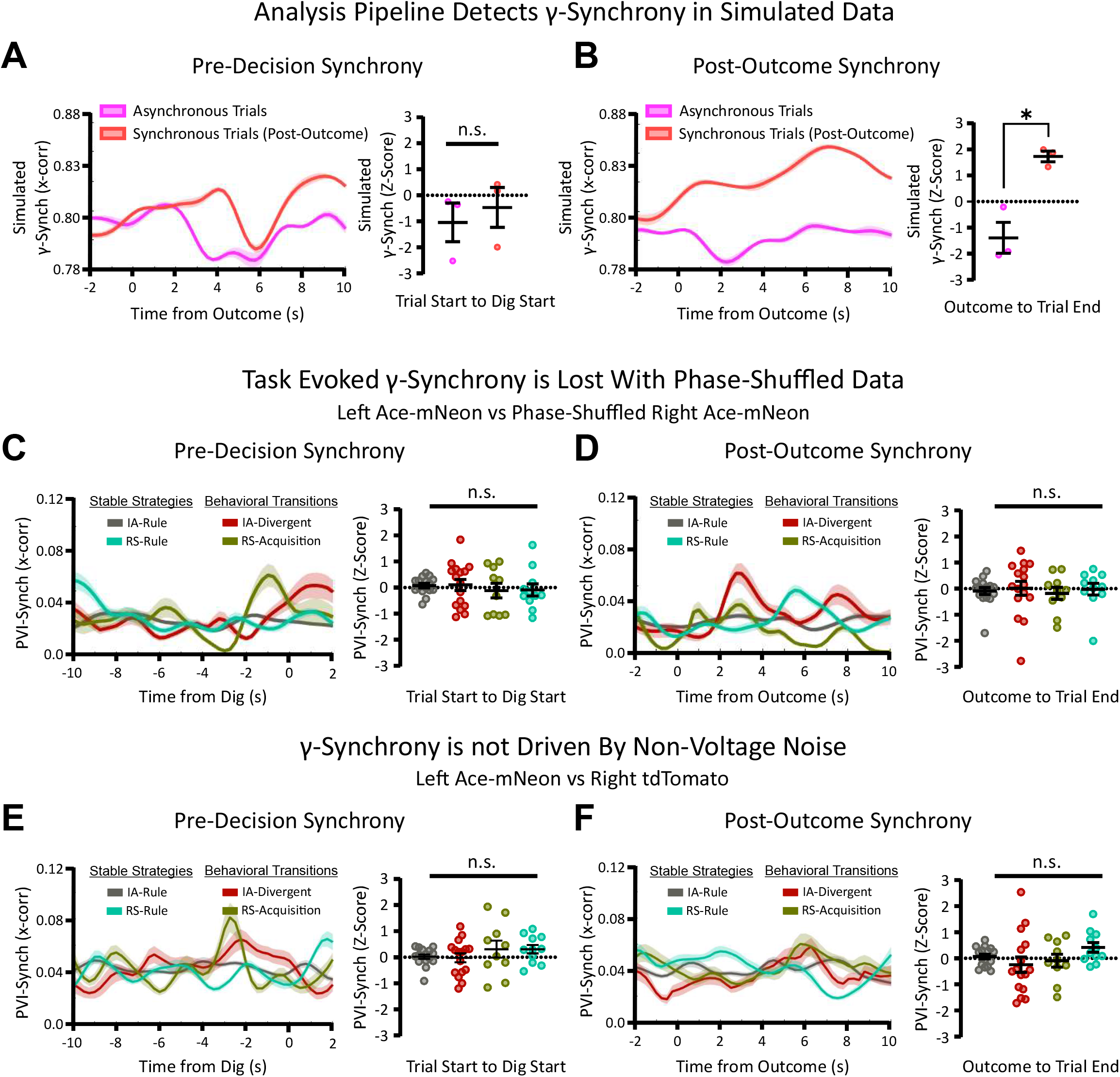
Validation of gamma synchrony analysis using simulated and phase-scrambled data. Validation methods demonstrate that our analysis pipeline accurately identifies phase-related synchrony in gamma-band activity. A-B) To confirm that our pipeline can detect time-resolved γ-Synchrony, we applied it to simulated data in which gamma oscillations were masked by realistic broadband, shared, and independent noise. We generated datasets for three ‘subjects,’ each containing two trials designated as ‘high’ in-phase synchrony trials during the post-outcome period (10 seconds between outcome and trial end), while all other epochs featured asynchronous gamma phases. The pipeline successfully identified enhanced γ-synchrony specifically during the post-outcome period of the synchronous trials (B). C–D) To ensure that observed synchrony in real data was not an artifact of the pipeline or driven by shared power or noise structure, we recalculated γ-synchrony on the PVI dataset after disrupting cross-hemispheric phase relationships using a Fourier-based phase-scrambling method. This manipulation abolished the increase in γ-Synchrony previously observed on IA-Divergent and RS-Acquisition trials (Fig. 2E vs. D), demonstrating that intact phase relationships are necessary for our pipeline to detect γ-Synchrony.

## Notes

### Competing Interest Statement

The authors have declared no competing interest.

### Summary of Updates

Added Figure captions which were missing from the previously uploaded version.

## REFERENCES

1. Green, M.F., Llerena, K., and Kern, R.S. (2015). The “Right Stuff” Revisited: What Have We Learned About the Determinants of Daily Functioning in Schizophrenia? Schizophrenia Bulletin 41, 781–785. 10.1093/schbul/sbv018.

2. Zhu, C., Kwok, N.T., Chan, T.C., Chan, G.H., and So, S.H. (2021). Inflexibility in Reasoning: Comparisons of Cognitive Flexibility, Explanatory Flexibility, and Belief Flexibility Between Schizophrenia and Major Depressive Disorder. Front. Psychiatry 11, 609569. 10.3389/fpsyt.2020.609569.

3. Boudewyn, M.A., and Carter, C.S. (2018). Evolving Concepts in Brain Oscillations and Cognitive Control in Schizophrenia. Biological Psychiatry 84, 632–633. 10.1016/j.biopsych.2018.08.017.

4. Collins, D.P., Anastasiades, P.G., Marlin, J.J., and Carter, A.G. (2018). Reciprocal Circuits Linking the Prefrontal Cortex with Dorsal and Ventral Thalamic Nuclei. Neuron 98, 366–379.e4. 10.1016/j.neuron.2018.03.024.

5. Parnaudeau, S., Bolkan, S.S., and Kellendonk, C. (2018). The Mediodorsal Thalamus: An Essential Partner of the Prefrontal Cortex for Cognition. Biological Psychiatry 83, 648–656. 10.1016/j.biopsych.2017.11.008.

6. Fornito, A., Harrison, B.J., Goodby, E., Dean, A., Ooi, C., Nathan, P.J., Lennox, B.R., Jones, P.B., Suckling, J., and Bullmore, E.T. (2013). Functional Dysconnectivity of Corticostriatal Circuitry as a Risk Phenotype for Psychosis. JAMA Psychiatry 70, 1143. 10.1001/jamapsychiatry.2013.1976.

7. Morris, R.W., Quail, S., Griffiths, K.R., Green, M.J., and Balleine, B.W. (2015). Corticostriatal Control of Goal-Directed Action Is Impaired in Schizophrenia. Biological Psychiatry 77, 187–195. 10.1016/j.biopsych.2014.06.005.

8. Boudewyn, M.A., Scangos, K., Ranganath, C., and Carter, C.S. (2020). Using prefrontal transcranial direct current stimulation (tDCS) to enhance proactive cognitive control in schizophrenia. Neuropsychopharmacology 45, 1877–1883. 10.1038/s41386-020-0750-8.

9. Sohal, V.S. (2022). Transforming Discoveries About Cortical Microcircuits and Gamma Oscillations Into New Treatments for Cognitive Deficits in Schizophrenia. American Journal of Psychiatry 179, 267–276. 10.1176/appi.ajp.20220147.

10. Strüber, D., and Herrmann, C.S. (2020). Modulation of gamma oscillations as a possible therapeutic tool for neuropsychiatric diseases: A review and perspective. International Journal of Psychophysiology 152, 15–25. 10.1016/j.ijpsycho.2020.03.003.

11. Adaikkan, C., and Tsai, L.-H. (2020). Gamma Entrainment: Impact on Neurocircuits, Glia, and Therapeutic Opportunities. Trends in Neurosciences 43, 24–41. 10.1016/j.tins.2019.11.001.

12. Fernandez-Ruiz, A., Sirota, A., Lopes-dos-Santos, V., and Dupret, D. (2023). Over and above frequency: Gamma oscillations as units of neural circuit operations. Neuron 111, 936–953. 10.1016/j.neuron.2023.02.026.

13. Horvath, A., Szucs, A., Csukly, G., Sakovics, A., Stefanics, G., and Kamondi, A. (2018). EEG and ERP biomarkers of Alzheimer’s disease: a critical review. Frontiers in Bioscience-Landmark 23, 183–220. 10.2741/4587.

14. Simmatis, L., Russo, E.E., Geraci, J., Harmsen, I.E., and Samuel, N. (2023). Technical and clinical considerations for electroencephalography-based biomarkers for major depressive disorder. npj Mental Health Research 2, 18. 10.1038/s44184-023-00038-7.

15. Sohal, V.S., Zhang, F., Yizhar, O., and Deisseroth, K. (2009). Parvalbumin neurons and gamma rhythms enhance cortical circuit performance. Nature 459, 698–702. 10.1038/nature07991.

16. Lewis, D.A., Curley, A.A., Glausier, J.R., and Volk, D.W. (2012). Cortical parvalbumin interneurons and cognitive dysfunction in schizophrenia. Trends in Neurosciences 35, 57–67. 10.1016/j.tins.2011.10.004.

17. Lodge, D.J., Behrens, M.M., and Grace, A.A. (2009). A Loss of Parvalbumin-Containing Interneurons Is Associated with Diminished Oscillatory Activity in an Animal Model of Schizophrenia. Journal of Neuroscience 29, 2344–2354. 10.1523/JNEUROSCI.5419-08.2009.

18. Cho, K.K.A., Hoch, R., Lee, A.T., Patel, T., Rubenstein, J.L.R., and Sohal, V.S. (2015). Gamma Rhythms Link Prefrontal Interneuron Dysfunction with Cognitive Inflexibility in Dlx5/6+/− Mice. Neuron 85, 1332–1343. 10.1016/j.neuron.2015.02.019.

19. Kujala, J., Jung, J., Bouvard, S., Lecaignard, F., Lothe, A., Bouet, R., Ciumas, C., Ryvlin, P., and Jerbi, K. (2015). Gamma oscillations in V1 are correlated with GABAA receptor density: A multi-modal MEG and Flumazenil-PET study. Sci Rep 5, 16347. 10.1038/srep16347.

20. Ferguson, B.R., and Gao, W.-J. (2018). PV Interneurons: Critical Regulators of E/I Balance for Prefrontal Cortex-Dependent Behavior and Psychiatric Disorders. Front. Neural Circuits 12, 37. 10.3389/fncir.2018.00037.

21. Murray, A.J., Woloszynowska-Fraser, M.U., Ansel-Bollepalli, L., Cole, K.L.H., Foggetti, A., Crouch, B., Riedel, G., and Wulff, P. (2015). Parvalbumin-positive interneurons of the prefrontal cortex support working memory and cognitive flexibility. Sci Rep 5, 16778. 10.1038/srep16778.

22. Cho, K.K.A., Davidson, T.J., Bouvier, G., Marshall, J.D., Schnitzer, M.J., and Sohal, V.S. (2020). Cross-hemispheric gamma synchrony between prefrontal parvalbumin interneurons supports behavioral adaptation during rule shift learning. Nature Neuroscience 23, 892–902. 10.1038/s41593-020-0647-1.

23. Spellman, T., Svei, M., Kaminsky, J., Manzano-Nieves, G., and Liston, C. (2021). Prefrontal deep projection neurons enable cognitive flexibility via persistent feedback monitoring. Cell 184, 2750–2766.e17. 10.1016/j.cell.2021.03.047.

24. Cho, K.K.A., Shi, J., Phensy, A.J., Turner, M.L., and Sohal, V.S. (2023). Long-range inhibition synchronizes and updates prefrontal task activity. Nature 617, 548–554. 10.1038/s41586-023-06012-9.

25. Bissonette, G.B., Powell, E.M., and Roesch, M.R. (2013). Neural structures underlying set-shifting: Roles of medial prefrontal cortex and anterior cingulate cortex. Behavioural Brain Research 250, 91–101. 10.1016/j.bbr.2013.04.037.

26. Marshall, J.D., Li, J.Z., Zhang, Y., Gong, Y., St-Pierre, F., Lin, M.Z., and Schnitzer, M.J. (2016). Cell-Type-Specific Optical Recording of Membrane Voltage Dynamics in Freely Moving Mice. Cell 167, 1650–1662.e15. 10.1016/j.cell.2016.11.021.

27. Fries, P. (2015). Rhythms for Cognition: Communication through Coherence. Neuron 88, 220–235. 10.1016/j.neuron.2015.09.034.

28. Buzsáki, G., and Wang, X.-J. (2012). Mechanisms of Gamma Oscillations. Annu. Rev. Neurosci. 35, 203–225. 10.1146/annurev-neuro-062111-150444.

29. Bolkan, S.S., Stujenske, J.M., Parnaudeau, S., Spellman, T.J., Rauffenbart, C., Abbas, A.I., Harris, A.Z., Gordon, J.A., and Kellendonk, C. (2017). Thalamic projections sustain prefrontal activity during working memory maintenance. Nature Neuroscience 20, 987–996. 10.1038/nn.4568.

30. Haziza, S., Chrapkiewicz, R., Zhang, Y., Kruzhilin, V., Li, J., Li, J., Delamare, G., Swanson, R., Buzsáki, G., Kannan, M., et al. (2025). Imaging high-frequency voltage dynamics in multiple neuron classes of behaving mammals. Cell, S0092867425007305. 10.1016/j.cell.2025.06.028.

31. Whittington, M.A., Traub, R.D., and Jefferys, J.G.R. (1995). Synchronized oscillations in interneuron networks driven by metabotropic glutamate receptor activation. Nature 373, 612–615. 10.1038/373612a0.

32. Fries, P. (2005). A mechanism for cognitive dynamics: neuronal communication through neuronal coherence. Trends in Cognitive Sciences 9, 474–480. 10.1016/j.tics.2005.08.011.

33. Tiesinga, P., and Sejnowski, T.J. (2009). Cortical Enlightenment: Are Attentional Gamma Oscillations Driven by ING or PING? Neuron 63, 727–732. 10.1016/j.neuron.2009.09.009.

34. Jackson, A.D., Cohen, J.L., Phensy, A.J., Chang, E.F., Dawes, H.E., and Sohal, V.S. (2024). Amygdala-hippocampus somatostatin interneuron beta-synchrony underlies a cross-species biomarker of emotional state. Neuron 112, 1182–1195.e5. 10.1016/j.neuron.2023.12.017.

35. Lourenço, J., Pacioni, S., Rebola, N., van Woerden, G.M., Marinelli, S., DiGregorio, D., and Bacci, A. (2014). Non-associative Potentiation of Perisomatic Inhibition Alters the Temporal Coding of Neocortical Layer 5 Pyramidal Neurons. PLoS Biol 12, e1001903.10.1371/journal.pbio.1001903.

36. Parnaudeau, S., O’Neill, P.-K., Bolkan, S.S., Ward, R.D., Abbas, A.I., Roth, B.L., Balsam, P.D., Gordon, J.A., and Kellendonk, C. (2013). Inhibition of Mediodorsal Thalamus Disrupts Thalamofrontal Connectivity and Cognition. Neuron 77, 1151–1162. 10.1016/j.neuron.2013.01.038.

37. Benoit, L.J., Canetta, S., and Kellendonk, C. (2022). Thalamocortical Development: A Neurodevelopmental Framework for Schizophrenia. Biological Psychiatry 92, 491–500. 10.1016/j.biopsych.2022.03.004.

38. Sohal, V.S. (2016). How Close Are We to Understanding What (if Anything) Oscillations Do in Cortical Circuits? Journal of Neuroscience 36, 10489–10495.10.1523/JNEUROSCI.0990-16.2016.

39. Cardin, J.A. (2016). Snapshots of the Brain in Action: Local Circuit Operations through the Lens of γ Oscillations. J. Neurosci. 36, 10496–10504. 10.1523/JNEUROSCI.1021-16.2016.

40. Zhu, X., Hagopian, L.L., Wallquist, K.E., and Sohal, V.S. (2025). Synaptic plasticity of prefrontal long-range inhibition regulates cognitive flexibility. Preprint at Cold Spring Harbor Laboratory, 10.1101/2025.06.27.662040.

41. Bosman, C.A., Lansink, C.S., and Pennartz, C.M.A. (2014). Functions of gamma-band synchronization in cognition: from single circuits to functional diversity across cortical and subcortical systems. Eur J of Neuroscience 39, 1982–1999. 10.1111/ejn.12606.

42. Canetta, S.E., Holt, E.S., Benoit, L.J., Teboul, E., Sahyoun, G.M., Ogden, R.T., Harris, A.Z., and Kellendonk, C. (2022). Mature parvalbumin interneuron function in prefrontal cortex requires activity during a postnatal sensitive period. eLife 11. 10.7554/elife.80324.

43. Gong, Y., Huang, C., Li, J.Z., Grewe, B.F., Zhang, Y., Eismann, S., and Schnitzer, M.J. (2015). High-speed recording of neural spikes in awake mice and flies with a fluorescent voltage sensor. Science 350, 1361–1366. 10.1126/science.aab0810.

44. Kannan, M., Vasan, G., Haziza, S., Huang, C., Chrapkiewicz, R., Luo, J., Cardin, J.A., Schnitzer, M.J., and Pieribone, V.A. (2022). Dual-polarity voltage imaging of the concurrent dynamics of multiple neuron types. Science 378. 10.1126/science.abm8797.

45. Chu, J., Oh, Y., Sens, A., Ataie, N., Dana, H., Macklin, J.J., Laviv, T., Welf, E.S., Dean, K.M., Zhang, F., et al. (2016). A bright cyan-excitable orange fluorescent protein facilitates dual-emission microscopy and enhances bioluminescence imaging in vivo. Nat Biotechnol 34, 760–767. 10.1038/nbt.3550.

46. Theiler, J., Eubank, S., Longtin, A., Galdrikian, B., and Doyne Farmer, J. (1992). Testing for nonlinearity in time series: the method of surrogate data. Physica D: Nonlinear Phenomena 58, 77–94. 10.1016/0167-2789(92)90102-s.

